# The human-specific miR-6762-5p is an activator of RhoA GTPase enhancing *Shigella flexneri* intercellular spreading

**DOI:** 10.1101/2022.11.28.518194

**Authors:** Caroline Reisacher, Estelle Saifi, Elisabeth Ageron-Ardila, Robert Theodor Mallmann, Norbert Klugbauer, David Skurnik, Laurence Arbibe

**Affiliations:** Université Paris Cité, INSERM, CNRS, Institut Necker Enfants Malades, F-75015 Paris, France; Service de microbiologie clinique, AP-HP, Hôpital Necker Enfants Malades, F-75015 Paris, France; Institut für Experimentelle und Klinische Pharmakologie und Toxikologie, Fakultät für Medizin, Albert-Ludwigs-Universität Freiburg, Albertstr. 25, 79104 Freiburg, Germany

## Abstract

MicroRNAs have recently emerged as major players in host-bacterial pathogens interaction, either as part of the host defense mechanism to neutralize infection or as a bacterial arsenal aimed at subverting host cell functions. Here we identify the newly evolutionary emerged human microRNA miR-6762-5p as a new player in the host-*Shigella* interplay. A microarray analysis in infected epithelial cells allowed the detection of this miRNA exclusively during the late phase of infection. Conditional expression of miR-6762-5p combined with a transcriptome analysis indicated a role in cytoskeleton remodeling. Likewise, miR-6762-5p enhanced stress fibers formation through RhoA activation and *in silico* analysis identified several regulators of RhoA activity as potential direct transcriptional targets. We further showed that miR-6762-5p expression induces an increase in *Shigella* intercellular spreading, while miR-6762-5p inhibition reduced bacterial dissemination. Overall, we have identified a human-specific miR-6762-5p acting specifically at the *Shigella* dissemination step. We propose a model in which the expression of miR-6762-5p induces cytoskeleton modifications through RhoA activation to achieve a successful dissemination of *Shigella* in the host.

## Introduction

*Shigella flexneri* (hereafter, *Shigella*) is a Gram-negative bacterium infecting the human recto-colic epithelium, causing bacillary dysentery. *Shigella*’s intracellular life cycle requires the control and remodeling of the host cytoskeleton, in particular the actin network (for review: Valencia-Gallardo et al., 2015). After entry and proliferation into the host cell, *Shigella* uses actin-based mobility (ABM), a mechanism by which the bacterium induces actin nucleation at one of its poles to create thrust, enabling the bacterial movement inside the host cytosol. This movement is directed towards the host cell membrane, to spread to the uninfected adjacent cell. The cell-to-cell spreading involves a reduction of cell-cell tension enabling the formation of a protrusion containing the bacterium, which is resolved by a clathrin-dependent process (Fukumatsu et al., 2012, Duncan-Lowey et al., 2020).

MicroRNAs (miRNAs) are small RNAs of about 22nt encoded into eukaryotic genomes. Their role in the post-transcriptional regulation of genes is being extensively studied. MiRNAs bind target mRNAs through partial complementarity and recruit molecular complexes inducing translational arrest, deadenylation, or degradation of the targeted mRNA. MiRNAs have been recognized for their role in the complex interplay between the host and bacterial pathogens (for review, Riahi Rad et al., 2021). In the context of *Shigella* infection, a microscopy-based high-throughput screening of a library of miRNA mimics has led to the identification of miRNAs regulating *Shigella* intra-cellular lifestyle (Sunkavalli et al., 2017; Aguilar et al., 2019). Likewise, the miR-29b-2-5p was shown to increase filopodia formation during the entry process by direct targeting of the Cdc42 and RhoF inhibitor UNC5C (Sunkavalli et al., 2017). This screening technique also enabled the characterization of 2 miRNAs impacting the ABM by targeting the N-WASP nucleator protein involved in actin polymerization, thereby decreasing bacterial intracellular mobility and dissemination (Aguilar et al., 2019). However to date, only a few miRNAs have been identified in the regulation of *Shigella* intra-cellular lifestyle.

Here, we provide a global overview of differentially expressed miRNAs in *Shigella*-infected colon carcinoma cells HCT116. Among those, we characterized the host human-specific miR-6762-5p for which the expression became detectable only at late infection time. We demonstrate that this miRNA induces the activation of the RhoA GTPase, thus promoting bacterial dissemination.

## Results

### MiR-6762-5p and its host gene TPCN1 are induced by *Shigella flexneri* infection

To provide a global view of the miRNAs differentially expressed during *Shigella* infection, we performed a microarray analysis of miRNA expression in the infected enterocytic HCT116 cells at 0.5- and 5-hours post-infection (hpi). This analysis uncovered a list of 32 mature miRNAs and 7 hairpins which were differentially regulated at 30 min post-infection, as compared to the non-infected samples. At 5 hpi, 30 mature miRNAs and 4 hairpins were differentially expressed (Figs 1A, 1B, S1 and Tables S1 & S2). Interestingly, among the differentially expressed miRNAs (DEM) expression of only 2 mature miRNAs (miR-4767 and miR-5100) and 1 hairpin (miR-5100) were commonly modified at both early and late infection time, highlighting a different pattern of the host miRNA response during early and late phases of infection.

**Figure 1.**
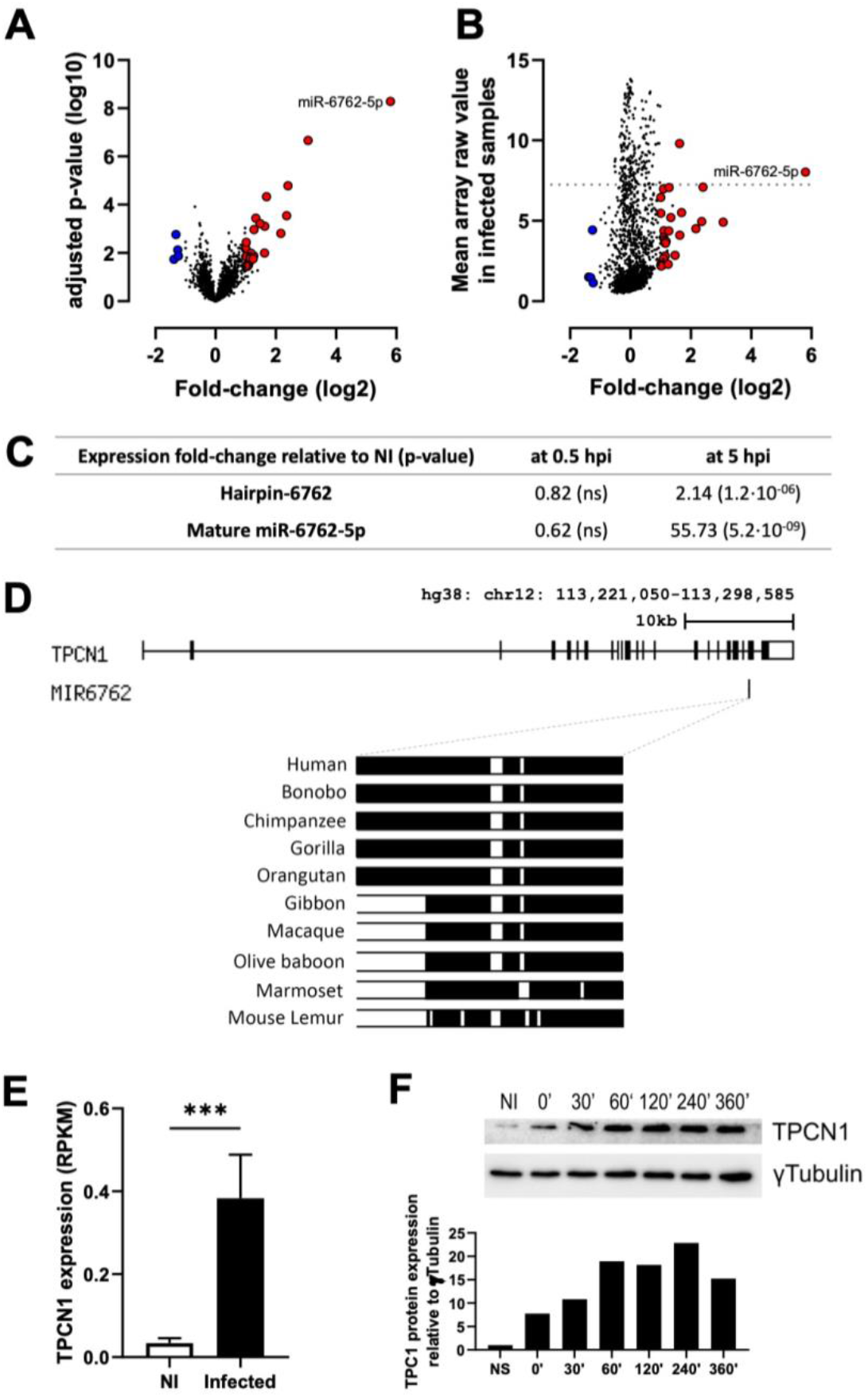
Global analysis of differentially expressed miRNAs in *Shigella flexneri* reveal the induction of miR-6762-5p. A- Volcano plot of miRNAs relative expression in *Shigella*-infected HCT116 cells 5 hpi compared to non-infected cells, expressed as log2(fold-change). In red are upregulated miRNAs, with higher than 2-fold-change expression and adjusted p-value under 0.05. In blue are downregulated miRNAs, with lower than 2-fold-change expression and adjusted p-value under 0.05. B- Plot representing the raw expression of miRNAs in 5h infected samples compared to their fold-change relative to the non-infected sample. Color code corresponds to panel A. Dotted line demarcates 200 most highly expressed miRNAs (above) in infected samples. C- Relative expression of miR-6762-5p and its precursor at 30 min or 5h post-infection (n=3). (A-C) Statistical comparisons were performed using two-way ANOVA. D- MIR6762 localization into one of TPCN1 introns and sequence comparison of human *MIR6762* gene locus with various monkeys. In black is represented aligned nucleotides and in white gaps in the alignment. E- mRNA expression of TPCN1 in non-infected and *Shigella*-infected HCT116 samples 5h hpi (n=3). Statistical comparison was performed using t-test. (***, p<0.001) F- Western blot analysis of TPCN1 protein expression during infection of HCT116 cells. Relative densitometry quantification is represented underneath. The immunoblot is representative of n=3 experiments.

Strikingly, miR-6762-5p was the most upregulated miRNAs with a 55-fold change at 5 hpi. This robust up-regulation was in part explained by a lack of miR-6762-5p detection above background signal in the non-infected samples. Based on raw microarray detection values, the expression of miR-6762-5p ranks among the 200 most expressed miRNAs in these cells at 5 hpi (Fig1B). Moreover, the miR-6762 precursor hairpin (pre-miR-6762) was also upregulated two-fold by *Shigella* infection. For both pre-miRNA and mature miR-6762-5p, the induction of expression was specifically detected at 5h thus at late phase of infection (Fig 1C). Evolutionary sequence analysis of miR-6762-5p revealed the sharing of this sequence only between human and evolutionary close apes (Bonobo, Chimpanzee, Gorilla, and Orangutan; Fig 1D), corresponding to the host specificity of *Shigella*. Additionally, the *MIR6762* gene is located at the 3’-end of an intron of the Two-Pore Channel 1 (*TPCN1* gene) and is classified as a mirtron (Ladewig et al., 2012; Wen et al., 2015; Fig 1D). As such, miR-6762-5p is expected to be co-expressed with the *TPCN1* transcript, after which the splicing of *TPCN1* mRNA liberates the intron bearing miR-6762-5p, further debranched and matured as a precursor. Here, we noted an increased expression of *TPCN1* transcript at 5 hpi and a gradual increase of TPC1 protein until 1 hpi after which the expression plateaued (Figs 1E and 1F). Thus, consistent with the gradual increase of the host gene *TPCN1* expression, our data identify the ape-specific host miR-6762-5p as a miRNA specifically induced upon late phase of *Shigella* infection.

### MiR-6762-5p induces the formation of stress fibers through RhoA activation

To unravel the role of the host miR-6762-5p, we stably transduced HCT116 and HeLa cells with a lentiviral Tet-On construction allowing a doxycycline-inducible expression of miR-6762-5p or a non-targeting (NTC) control miRNA (hereafter noted: HCT116^miR^, HCT116^NTC^, HeLa^miR^, and HeLa^NTC^; Fig 2A). Although a low basal transgene expression (i.e. without adding doxycycline) was detected by RT-qPCR (Fig 2B), the system was readily inducible, with increased miR-6762-5p expression by about 12 and 100-fold in the HCT116^miR^ and HeLa^miR^ cells respectively, at 48 hours post-doxycycline administration (Fig 2B). To determine the target pathways of miR-6762-5p, we further analyzed gene expression changes by high-throughput RNA sequencing (Fig 2C). By comparing doxycycline-induced miR-6762-5p and NTC cell lines, we identified the 10 most significant gene ontologies pathways enriched upon miR-6762-5p expression (Fig 2C, Tables S3-S6). Remarkably, these pathways pointed towards cytoskeleton remodeling in both cell lines. We confirmed the differential gene expression of some cytoskeleton-related genes by RT-qPCR, as shown by increased detection of MYO1E and FBLIM1 transcripts in both HCT116 and HeLa cell lines (Fig S2). Moreover, specifically in HCT116, we observed a potent increase in TMSB4X, an actin sequestering protein, expression upon miR-6762-5p induction (Safer et al., 1991). Likewise, to define a functional impact on actin polymerization, we performed phalloidin staining of F-actin and found that induction of miR-6762-5p expression led to a significant increase of actin stress fibers in HeLa^miR^ cells (Fig 2D). These observations led us to investigate whether miR-6762-5p expression led to the activation of small GTPases, as these are major modulators of cytoskeletal functions. Likewise, miR-6762-5p expression consistently increased RhoA activation in both HCT116 and HeLa cells (Fig 3A), while the activations of Rac1 and CDC42 were not impacted (Fig S4). Importantly, transfection of miR-6762-5p inhibitor or addition of Y27632 - a ROCK inhibitor acting downstream of RhoA, reversed stress fibers formation in HeLa cells (Fig 3B-C). Similar experiments performed in the HCT116 cells showed that pharmacological inhibition of RhoA or use of the miR-6762-5p inhibitor both led to a increase in filopodia extensions, suggesting that, as reported in neurons (Yuan et al. 2002), RhoA activity in this cell line triggered a cytoskeletal reorganization inhibiting filopodia extension (Fig S3).

**Figure 2.**
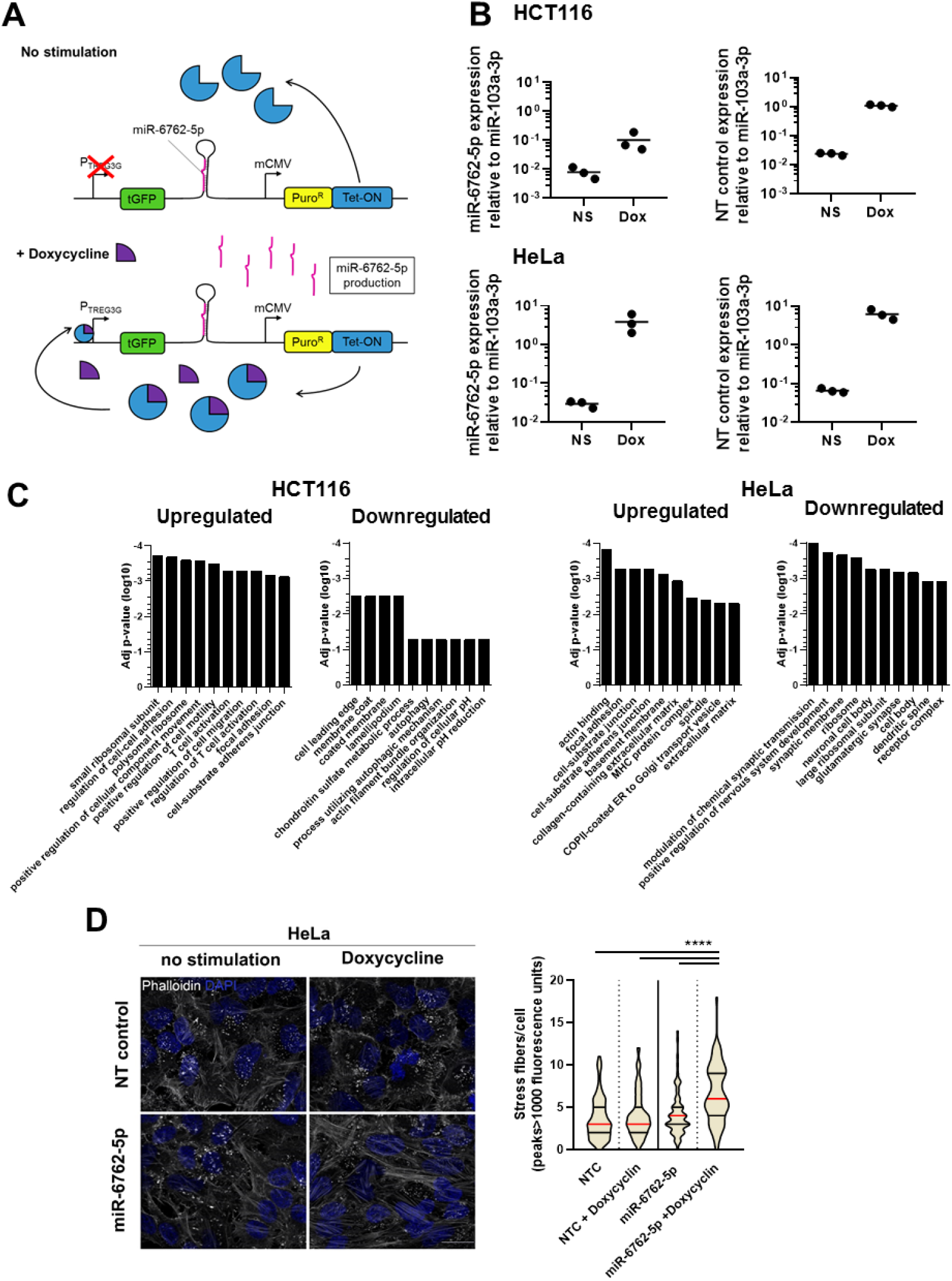
miR-6762-5p targets the cytoskeleton regulation in HCT116 and HeLa cells. A- Scheme of the inducible miRNAs expressing system. B- Validation by stem-loop RT-qPCR of inducible expression of miR-6762-5p and non-targeting control miRNAs in HCT116 and HeLa transduced cells. C- Top 10 most enriched GO pathways in upregulated and downregulated genes in HCT116 and HeLa cells, sorted by adjusted p-values. Gene expression quantified by RNA sequencing experiments of *n*=3 separate experiments. D- Fluorescent labeling of F-Actin with Phalloidin (white) and DAPI labeling of DNA (blue) in HeLa^miR^ and HeLa^NTC^ with and without doxycycline induction of miRNAs expression. Quantification of minimum 75 cells in 5 pictures of actin stress fibers per cells. Data are representative of *n*=3 experiments. Scale: 20µm. Statistical comparisons were performed using one-way ANOVA. (****, p<0.0001)

**Figure 3.**
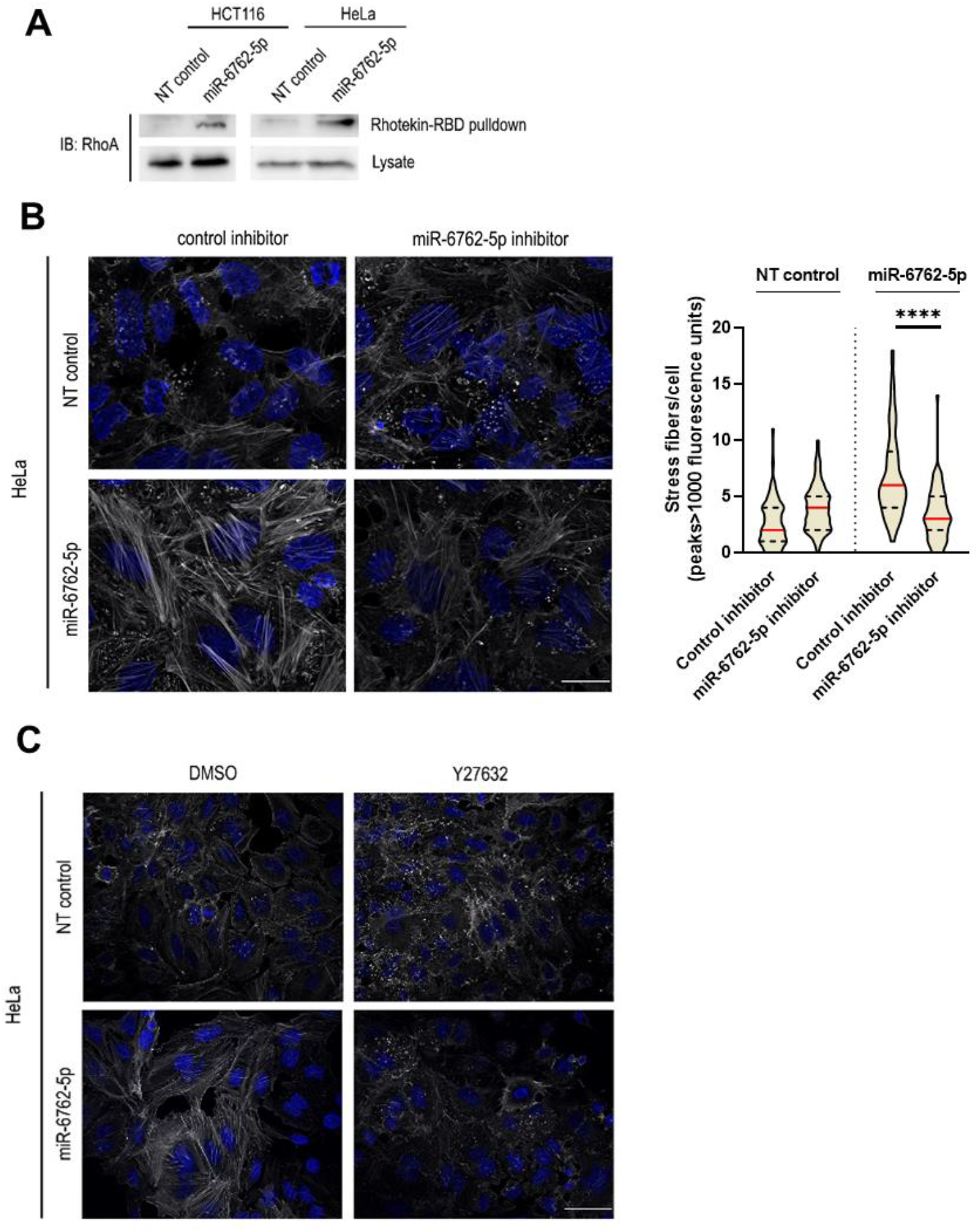
miR-6762-5p promotes RhoA activation during late infection. A- RhoA activation assay by GTP-bound RhoA pulldown followed by Western blot reative quantification. Comparison of RhoA activation in induced HCT116^NTC^ vs. HCT116^miR^ and HeLa^NTC^ vs. HeLa^miR^. Total lysate Western blot for relative quantification. Representative images of n=2 experiments. B- Fluorescent labeling of F- Actin with Phalloidin (white) and DAPI labeling of DNA (blue) in HCT116 and HeLa with doxycycline induction of miRNA expression, transfected with a control miRNA inhibitor vs. a miR-6762-5p inhibitor. Data are representative of *n*=2 experiments. Scale: 20µm. Representative images of n= 2 experiments. Statistical comparisons were performed using one-way ANOVA. (****, p<0.0001) C- Fluorescent labeling of F-Actin with Phalloidin (white) and DAPI labeling of DNA (blue) in HeLa^miR^ and HeLa^NTC^ with doxycycline induction of miRNAs expression treated with ROCK inhibitor 15µM Y27623 or carrier DMSO for 1h. Statistical significance was assessed using one-way ANOVA with mean comparisons two by two and p-value adjustments for multiple testing. Representative images of n= 2 experiments.

To investigate potential mechanisms by which miR-6762-5p expression modulated RhoA activity, we analyzed from the RNA-seq data the expression level of the main RhoA regulators, namely the RhoA guanine nucleotide exchange factors (GEFs), the GTPases-activating proteins (GAPs), and GDP dissociation inhibitors (GDIs). GAPs enable the GTP hydrolysis of the rhoGTPases, thus rendering them inactive. Interestingly, in HeLa cells, miR-6762-5p induced a potent transcriptional downregulation of some members of the *ARHGAP*, a family of genes encoding RhoGAPs (Fig 4A). Remarkably, *in silico* analysis aimed at investigating a direct targeting of these transcripts by miR-6762-5p showed that all transcripts possessed at least one potential target site (Table 1). By contrast, miR-6762-5p-expressing HCT116 showed little modifications of these effectors (Fig 4B).

**Table 1:**
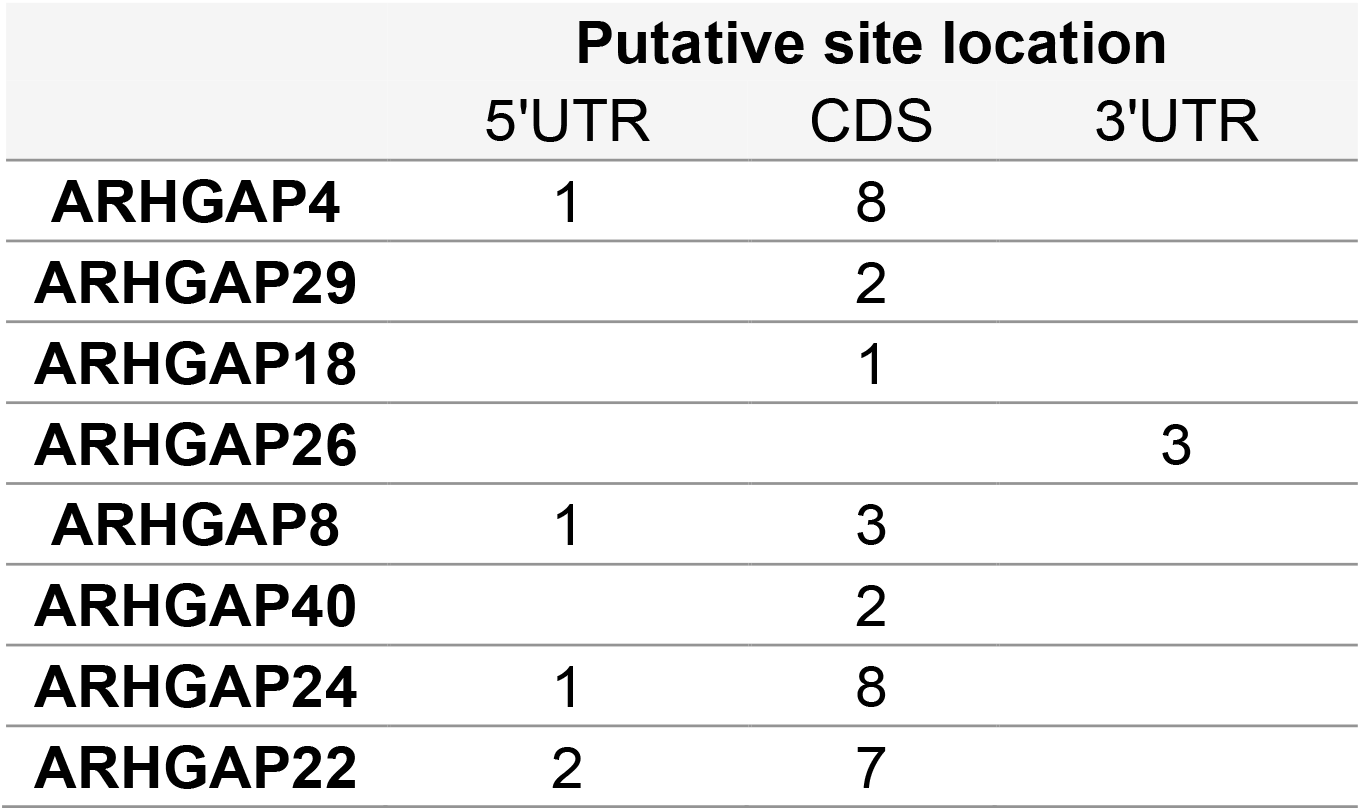
*In sillico* analysis of putative miR-6762-5p target sites on downregulated rhoGAP members. Number and location of putative miR-6762-5p target sites on ARHGAPs’ transcripts as identified by miRWalk algorithm.

|  | Putative site location |  |  |
| --- | --- | --- | --- |
|  | 5'UTR | CDS | 3'UTR |
| **ARHGAP4** | 1 | 8 |  |
| **ARHGAP29** |  | 2 |  |
| **ARHGAP18** |  | 1 |  |
| **ARHGAP26** |  |  | 3 |
| **ARHGAP8** | 1 | 3 |  |
| **ARHGAP40** |  | 2 |  |
| **ARHGAP24** | 1 | 8 |  |
| **ARHGAP22** | 2 | 7 |  |

**Figure 4.**
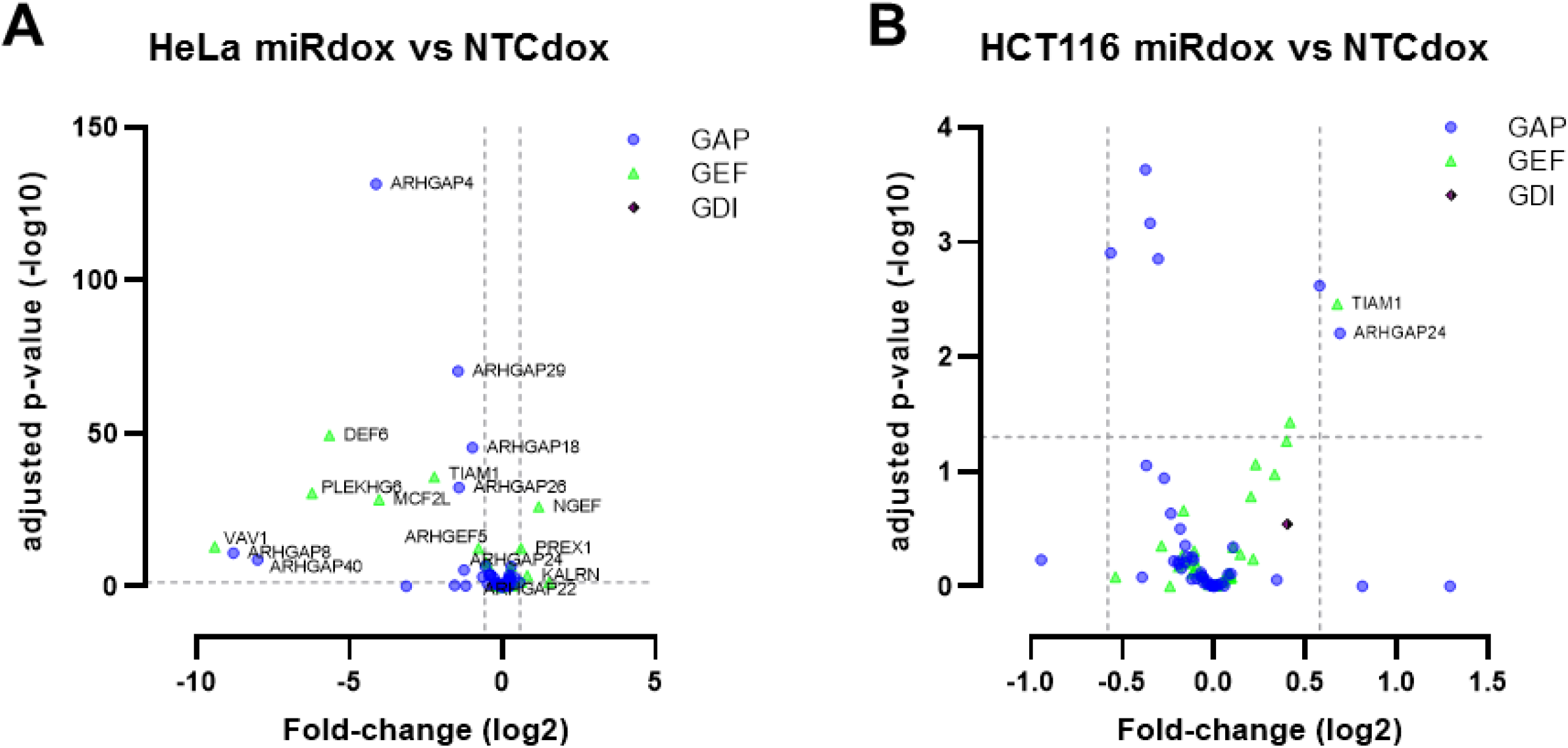
Analysis of RhoA activation regulators expression upon miR-6762-5p expression. A- Volcano plot of RhoA GAPs, GEFs, and GDIs representing their log2(Fold-Change) expression as function of their adjusted p-values in doxycyclin-induced HeLa^miR^ vs HeLa^NTC^. B- Volcano plot of RhoA GAPs, GEFs, and GDIs representing their log2(Fold-Change) expression as function of their adjusted p-values in doxicyclin-induced HCT116^miR^ vs HCT116^NTC^. Horizontal dotted line represents the threshold for p=0,05. Statistically significant values are represented above the line. Vertical dotted lines represent the fold-change threshold we selected for differentially expressed genes (left line: fold-change = −1,5; right line: fold-change = 1,5). Downregulated genes are thus represented left of the left line and upregulated genes, right of the right line. Statistical comparisons were performed using two-way ANOVA.

Overall, these data showed that miR-6762-5p remodels the host cytoskeleton through a selective activation of the RhoA GTPase.

### MiR-6762-5p expression promotes *Shigella* dissemination

Considering the impact of miR-6762-5p on RhoA activation and more generally on the host cytoskeleton, we reasoned that it should impact *Shigella*’s intracellular life cycle. We first confirmed that *Shigella* infection modulated RhoA activation, with an early and transient activation peak followed by a second wave of RhoA activation (Fig 5A). Likewise, phosphorylation of MLC2, a distal target of RhoA signaling, occurs at late phase of infection (Fig 5B). To investigate the effect of miR-6762-5p expression on the number of intracellular *Shigella* colonies, we performed gentamicin protection assay at early and late phases of infection in both HCT116^miR^ and HCT116^NTC^. The results showed that miRNA overexpression did not affect the number of intracellular bacteria at both infection times, thus suggesting no impact on the entry process and on bacterial proliferation within the target cell (Fig 6A-B). To assess the extent of intercellular spread of bacteria, we performed plaque formation assay by performing infection in confluent miR-induced HCT116 cells. MiR-6762-5p expression led to a significant increase of the average size of plaques, while not affecting their number (Fig 6C and data not shown). Importantly, plaque formation assay performed in HCT116 WT cells showed that the miR-6762-5p inhibitor conversely reduced the average size of the plaque (Fig 6D). Thus, blocking endogenous miR-6762-5p activity restricts *Shigella flexneri* dissemination. Overall, these results show that the production of miR-6762-5p during infection promotes bacterial dissemination without impacting bacterial entry or intracellular proliferation.

**Figure 5.**
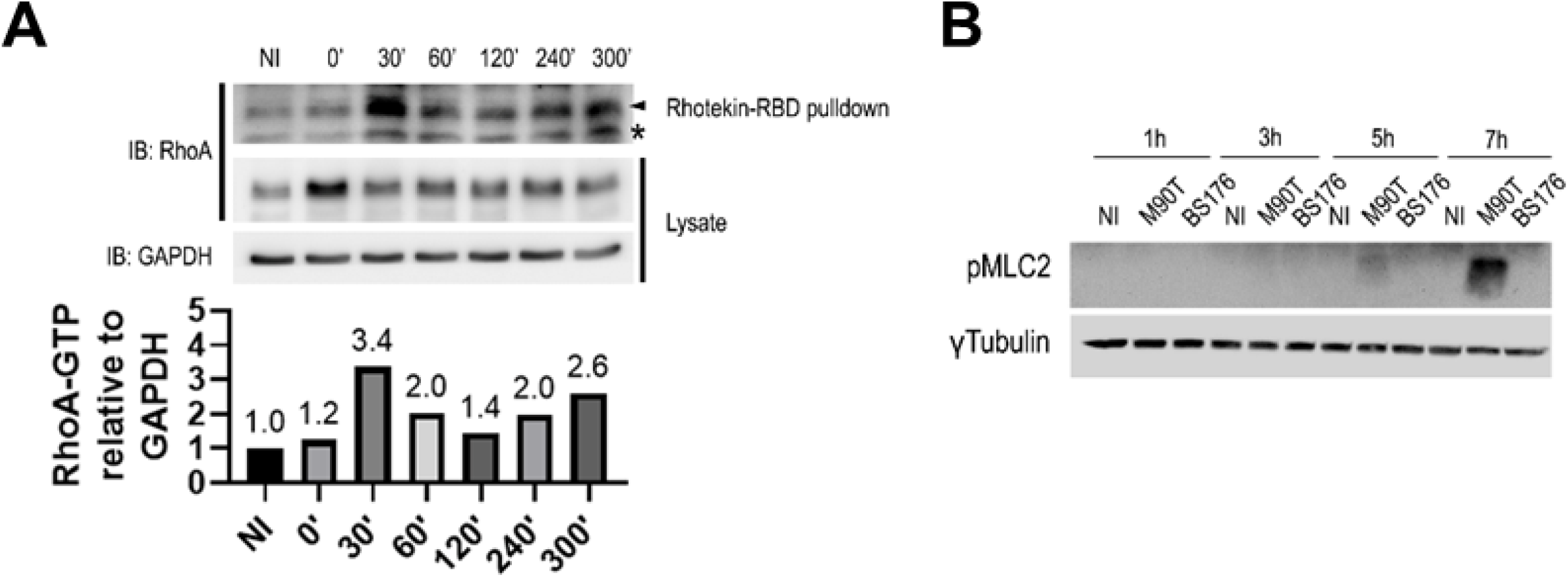
*Shigella* infection induces an early and late activation peak of RhoA. A- RhoA activation assay in *Shigella* infected wild-type HCT116 cells. Quantification corresponds to the ratio of active RhoA protein over GAPDH protein, determined by densitometry analysis. Representative image of n=2 independent experiments. B- Western blot analysis of MLC2 phosphorylation during infection. Representative image of n=2 independent experiments.

**Figure 6.**
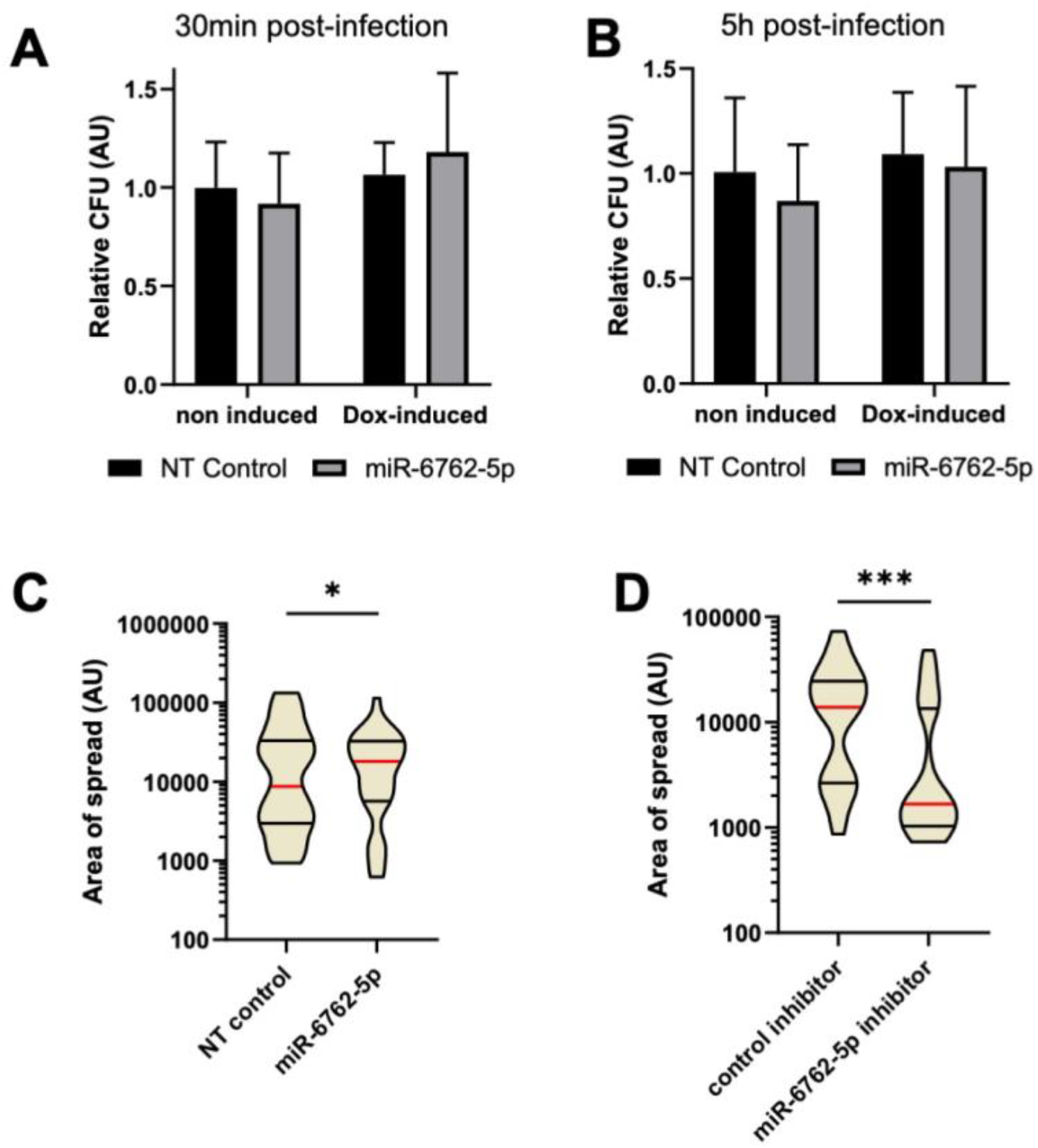
miR-6762-5p promotes bacterial spreading without impacting bacterial entry or intracellular proliferation. A- Gentamycin protection assay in HCT116^NTC^ vs HCT116^miR^ cells previously induced or not. Infection for 30min by *Shigella* at MOI 100. B- Gentamycin protection assay in HCT116^NTC^ vs HCT116^miR^ cells previously induced or not. Infection for 5h by *Shigella* at MOI 100. C- Plaque assay comparing bacterial spreading area in confluent induced HCT116^NTC^ vs HCT116^miR^ cells. Infection for 24h by *Shigella* at MOI 0.01. Quantification of minimum *n*=30 plaques of 2 independent experiments. Statistical significance was assessed using t-test. D- Plaque assay comparing bacterial spreading area in confluent wild-type HCT116 cells previously transfected with control or miR-6762-5p inhibitor. Infection for 24h by *Shigella* at MOI 0.01. Quantification of minimum *n*=30 plaques of 2 independent experiments. Statistical significance was assessed using t-test. (*, p<0.05; ***, p<0.001)

## Discussion

In this work, we have provided a dynamic overview of differentially expressed miRNAs during the early and late steps of *Shigella* infection *in vitro*. These steps represent 2 stages of *Shigella*’s intracellular lifestyle: the entry phase, at which the bacterium enters in its vacuole of phagocytosis and then disrupts it to freely proliferate into the host cytoplasm; and the proliferation/dissemination phase (for review: Mellouk & Enninga, 2016; Nasser et al., 2022). These different steps of the bacterium’s lifestyle elicit distinct miRNA host response. Indeed, we could only identify 2 mature miRNAs that were commonly expressed 30 min and 5 h post-infection.

MiR-6762-5p was the most differentially expressed miRNAs in our analysis. No basal expression could be detected, and early infection did not elicit its expression. However, during late infection, its expression was amongst the 200 highest expressed miRNAs. Interestingly, miR-6762-5p is evolutionary new in the human genome as its gene is only shared with apes. Remarkably, *Shigella* is host-adapted to humans and non-human primates (i.e. apes) while the determinant of this host specificity is yet to be uncovered. As such, the matching specificity of this miRNA and *Shigella*’s host specificity is intriguing. Interestingly, miR-6762-5p is a putative mirtron located on the *TPCN1* gene, and we showed that the corresponding protein product TPC1 is increased upon *Shigella* infection. TPC1 is an endosomal ion channel, with reported functions in bacterial toxins uptake deserving further exploration in the context of *Shigella* infection (Davis et al., 2020; Castonguay et al., 2017). Nevertheless, we found that transcriptional induction of *TPCN1* correlated with the increase of miR-6762-5p expression, providing a plausible mechanism for triggering the expression of the mirtron. Mechanistically, mirtron production is known to result from the activity of mirtron-specific maturation effectors, namely the splicing machinery and the debranching enzyme DBR1. Likewise, the first step of mirtron maturation involves splicing of the host transcript allowing the release of the miRNA-containing intron. After debranching of the intron, the pre-miRNA bypasses the microprocessor and is directly exported out of the nucleus where it follows the canonical maturation route (Ruby et al., 2007). Thus, any modification in mirtron-processing effectors activity, or in the intron degradation pathway may account for the induction of miR-6762-5p expression during *Shigella* infection.

The development of conditionally inducible cell lines expressing miR-6762-5p combined with a transcriptome analysis led us to identify the actin cytoskeleton as a targeted pathway by miR-6762-5p. In this analysis, miR-6762-5p expression induces the formation of stress fibers through RhoA activation while the key miRNA targets sustaining this functional effect remain to be identified. RhoA activation can ensue from an increase in Rho activators – RhoGEFs or a decrease in Rho activity inhibitors – RhoGDIs sequesters and RhoGAP hydrolyzing enzymes. Based on our transcriptome analysis, we identified a small group of up-regulated GEFs and down-regulated GAPs. MiRNAs are mostly known for inducing translation arrest or degradation of an mRNA through partial sequence homology. Providing miR-6762-5p is acting on RhoA activation in a canonical microRNA function, the direct repression of one or more RhoGDIs or RhoGAP would explain RhoA activation. Following the previous assumption, *in silico* analysis identified 8 ARHGAPs with one or more putative target site for miR-6762-5p, associated to an expression negatively correlated to that of miR-6762-5p. These genes may be direct targets of miR-6762-5p. Transcriptional activation of target genes by microRNAs has also, though rarely, been reported (for review: Stavast & Erkeland, 2019). This non-canonical miRNA activity may be mediated through antisense binding of the microRNA to the target’s promoter sequence in the nucleus, leading to the recruitment of the transcription machinery to regulate transcription. Thus, RhoGEFs with expression correlated to miR-6762-5p may also, be direct targets accounting for the increased RhoA activity.

RhoGTPases are known to be activated during *Shigella* invasion of host cells. Type III secretion effectors specifically target the activation of Rac1, CDC42 and RhoA to promote actin remodeling, filopodia formation, and allow the engulfment of the bacterium inside the host cell (Mounier et al., 1999). Three main effectors are responsible for rhoGTPases activation: IpgB1 and IpaC activate Rac1 and CDC42, while IpgB2 acts as RhoGEF mimic to activate RhoA (Tran Van Nhieu et al., 1999; Alto et al., 2006). Our analysis identified 2 waves of RhoA activation during *Shigella* infection *in cellulo*, the late activation questioning a possible role in bacterial spreading. Efficient cell-to-cell spread implies that *Shigella* pushes outwardly against the plasma membrane, creating a membrane-bound cell extension (“protrusion”) that must project into adjacent cells. Studies focusing on the role of myosin II light chain (MLC) phosphorylation have clearly shown the importance of the actin–myosin II networks during *Shigella* intercellular spreading (Rathman et al., 2000; Lum & Morona, 2014). RhoA is a notorious trigger for MLC phosphorylation and activity (Maekawa et al., 1999; Amano et al., 1996). Thus, RhoA activity may contribute to bacterial spreading in the context of *Shigella* infection.

In conclusion, we have identified a human-specific miR-6762-5p expressed and acting specifically at the *Shigella* dissemination step. We thus propose a model in which the expression of miR-6762-5p induces cytoskeleton modifications through RhoA activation to achieve successful dissemination of *Shigella* in the host.

---

**Figure S1.**
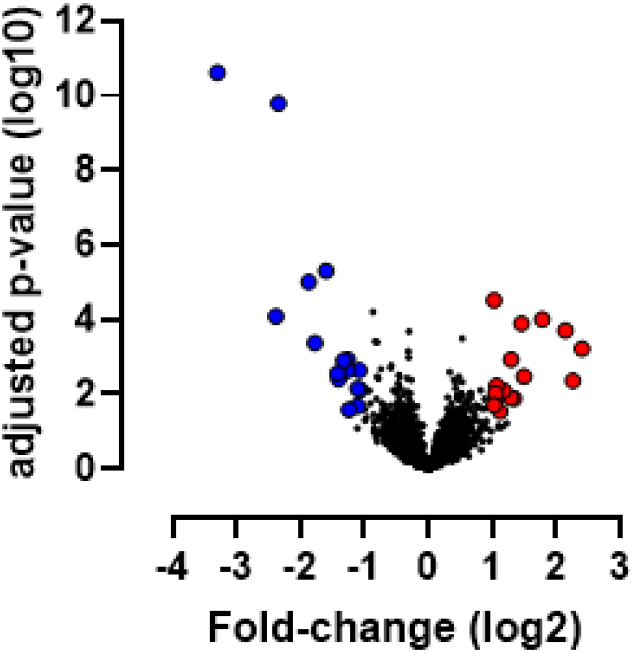
Global analysis of miRNAs expression changes at early infection time. Volcano plot of miRNAs relative expression in *Shigella*-infected HCT116 cells 0.5 hpi compared to non-infected cells, expressed as log2(fold-change). In red are upregulated miRNAs, with higher than 2-fold-change expression and adjusted p-value under 0.05. In blue are downregulated miRNAs, with lower than 2-fold-change expression and adjusted p-value under 0.05. Statistical comparisons were performed using two-way ANOVA.

**Figure S2.**
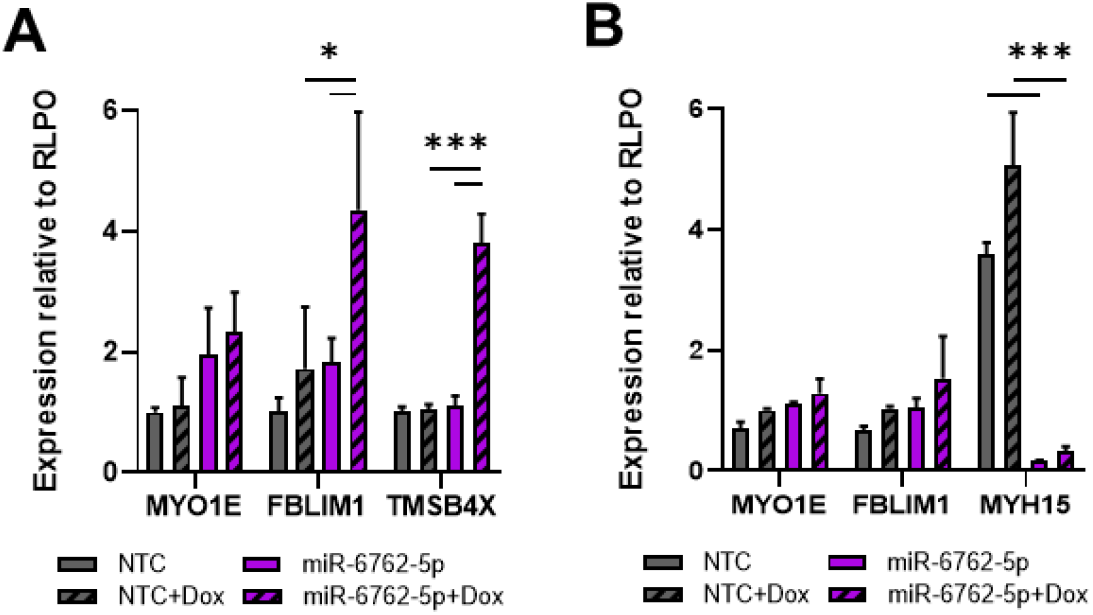
Validation of some differentially expressed cytoskeleton genes. A- Analysis by RT-qPCR of differentially expressed cytoskeletal genes (*MYO1E, FBLIM1, TMSB4X*) in HCT116^NTC^ vs. HCT116^miR^, induced vs non-induced. B- Analysis by RT-qPCR of differentially expressed cytoskeletal genes (*MYO1E, FBLIM1, MYH15*) in HeLa^NTC^ vs. HeLa^miR^, induced vs non-induced. Mean relative expression and SD from 3 independent experiments. Statistical comparisons were performed using two-way ANOVA. (*, p<0.05; ***, p<0.001)

**Figure S3.**
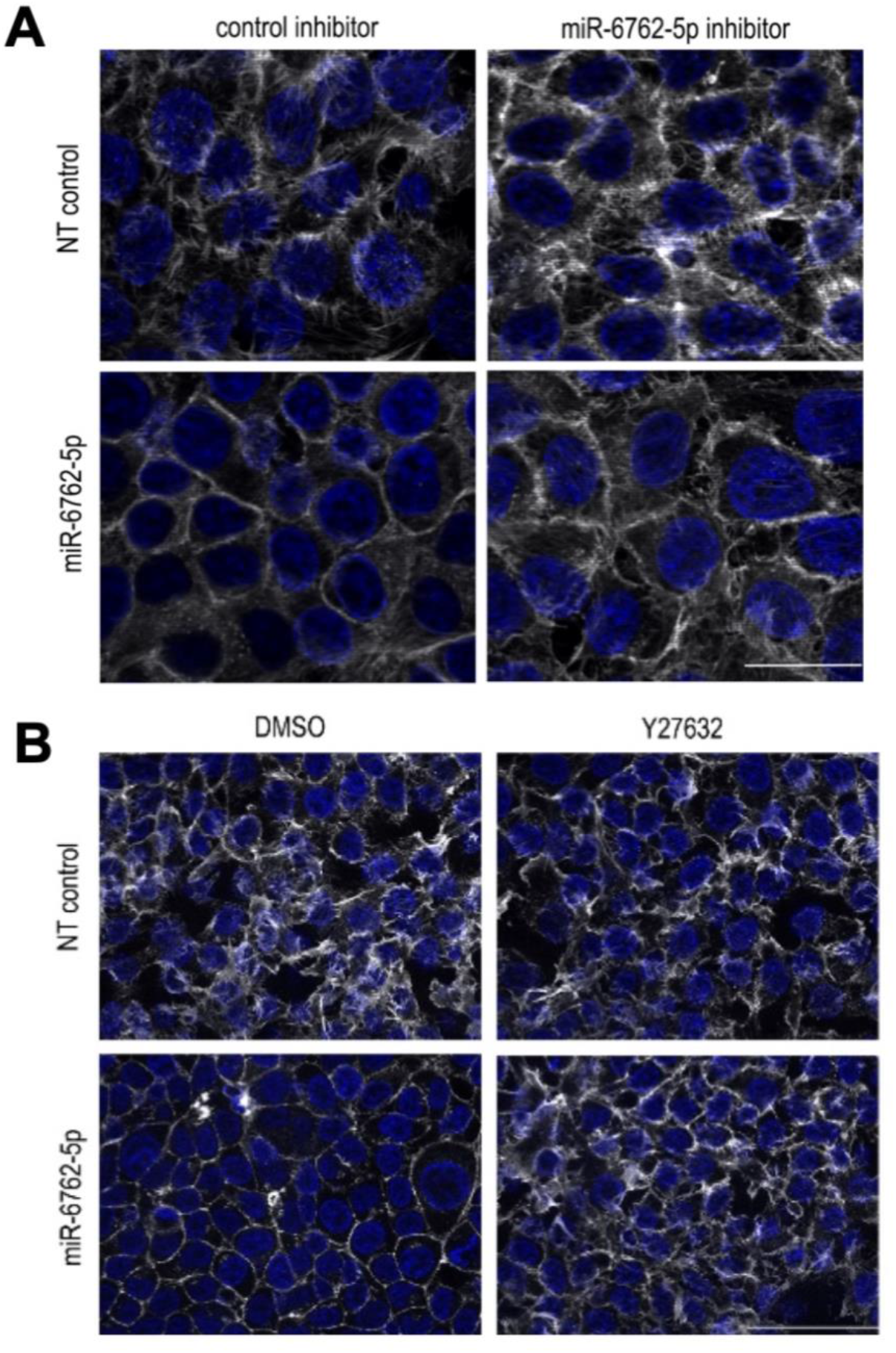
miR-6762-5p expression induces reduction of filopodia through ROCK. A - Fluorescent labeling of F-Actin with Phalloidin (white) and DAPI labeling of DNA (blue) in HCT116^miR^ and HCT116^NTC^ with doxycycline induction of miRNAs expression transfected with miR-6762-5p inhibitor or control miRNA inhibitor. Scale: 20µm. Representative images of n=2 experiments. B- Fluorescent labeling of F-Actin with Phalloidin (white) and DAPI labeling of DNA (blue) in HCT116^miR^ and HCT116^NTC^ with doxycycline induction of miRNAs expression treated with ROCK inhibitor 15µM Y27632 or carrier DMSO for 1h. Representative images of n=2 experiments.

**Figure S4.**
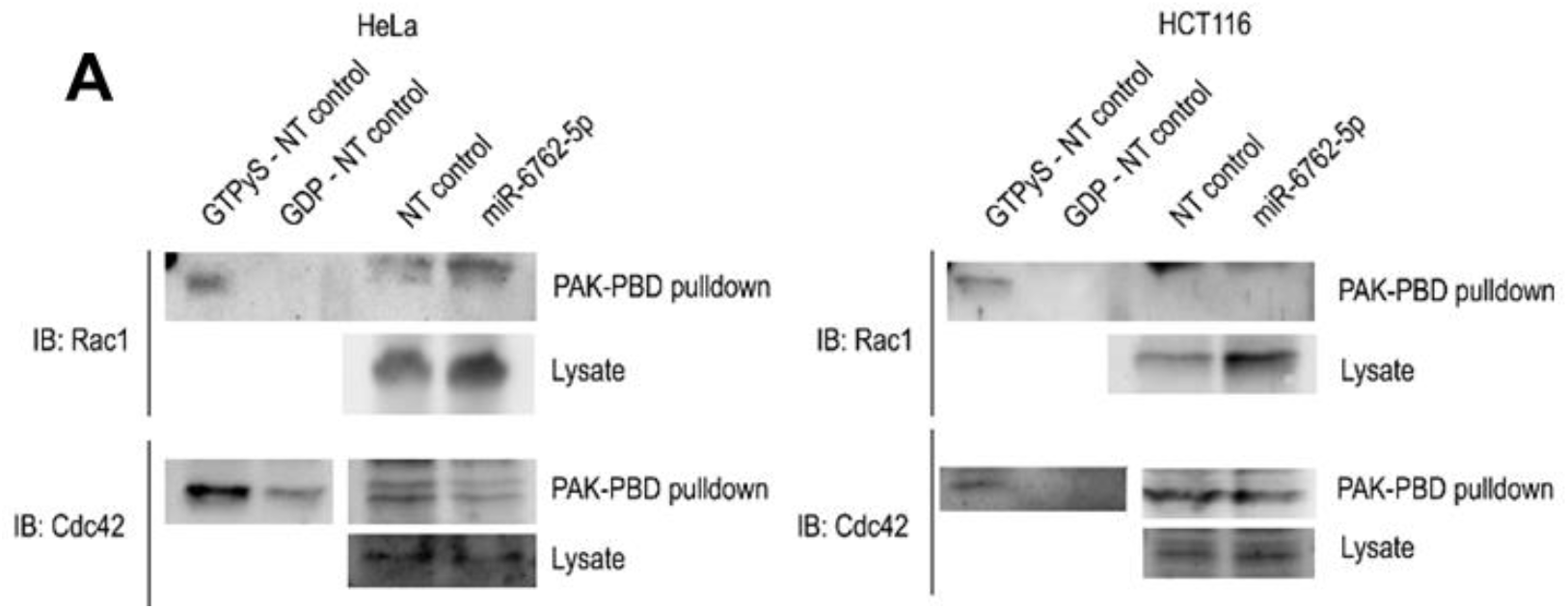
miR-6762-5p does not impact Rac1 or Cdc42 activation. Rac1 and Cdc42 activation assay by GTP-bound Rac1 and GTP-bound Cdc42 pulldown followed by Western blot. Total lysate Western blot for relative quantification. Comparison of induced HCT116^NTC^ vs. HCT116^miR^ and HeLa^NTC^ vs. HeLa^miR^. Representative images of n=2 experiments.

**Table S1:** microarray analysis of differentially regulated microRNAs at 0.5hpi.

| **Column ID** | **p-value(Shigella-Infected-30min vs. NI)** | **Fold-Change(Shigella-Infected-30min vs. NI)** |
| --- | --- | --- |
| MATURE_hsa-mir-27a-5p_at | 2,38913E-11 | -9,8329 |
| MATURE_hsa-mir-23a-5p_at | 1,58615E-10 | -5,07636 |
| MATURE_hsa-mir-4443_at | 4,97471E-06 | -3,02207 |
| MATURE_hsa-mir-424-3p_at | 9,89528E-06 | -3,65831 |
| HAIRPIN_hsa-mir-424_at | 1,85893E-05 | -2,29605 |
| MATURE_hsa-mir-200c-5p_at | 3,07025E-05 | 2,03698 |
| HAIRPIN_hsa-mir-4443_at | 3,60204E-05 | -2,47569 |
| MATURE_hsa-mir-4485-3p_at | 8,16867E-05 | -5,21852 |
| MATURE_hsa-mir-6779-5p_at | 9,97579E-05 | 3,423 |
| MATURE_hsa-mir-1229-5p_at | 0,000128328 | 2,73666 |
| MATURE_hsa-mir-1972_at | 0,00019669 | 4,40679 |
| HAIRPIN_hsa-mir-4485_at | 0,000207762 | -3,91959 |
| MATURE_hsa-mir-4440_at | 0,000425968 | -3,42326 |
| <u>MATURE_hsa-mir-5100_at</u> | 0,0006068 | 5,28934 |
| <u>HAIRPIN_hsa-mir-5100_at</u> | 0,000631503 | 4,07442 |
| MATURE_hsa-mir-3928-3p_at | 0,00116981 | -2,3994 |
| MATURE_hsa-mir-6833-5p_at | 0,00117762 | 2,44911 |
| MATURE_hsa-mir-4486_at | 0,00127444 | -2,50373 |
| HAIRPIN_hsa-mir-7975_at | 0,00221891 | 2,38591 |
| <u>MATURE_hsa-mir-4767_at</u> | 0,00227645 | -2,40246 |
| MATURE_hsa-mir-4532_at | 0,00230144 | -2,11031 |
| MATURE_hsa-mir-5580-3p_at | 0,00271767 | -2,66736 |
| HAIRPIN_hsa-mir-4454_at | 0,00304697 | 2,72624 |
| MATURE_hsa-mir-4454_at | 0,0033593 | 2,81454 |
| MATURE_hsa-mir-21-3p_at | 0,00393275 | -2,64127 |
| MATURE_hsa-mir-7975_at | 0,00433531 | 4,78462 |
| MATURE_hsa-mir-4684-3p_at | 0,00585758 | 2,08952 |
| MATURE_hsa-mir-192-5p_at | 0,0070612 | -2,14843 |
| MATURE_hsa-mir-1909-3p_at | 0,00764481 | -2,09134 |
| MATURE_hsa-mir-4444_at | 0,00802623 | 2,26243 |
| HAIRPIN_hsa-mir-7977_at | 0,00871132 | 2,22839 |
| MATURE_hsa-mir-4793-5p_at | 0,00936488 | 2,06102 |
| MATURE_hsa-mir-7977_at | 0,012525 | 2,45237 |
| MATURE_hsa-mir-6074_at | 0,0129759 | 2,52934 |
| MATURE_hsa-mir-5739_at | 0,0141426 | 2,0674 |
| MATURE_hsa-mir-34a-3p_at | 0,0196214 | 2,04108 |
| MATURE_hsa-mir-6824-5p_at | 0,0208718 | -2,14906 |
| MATURE_hsa-mir-584-3p_at | 0,0259074 | -2,36536 |
| MATURE_hsa-mir-451a_at | 0,0292801 | 2,16923 |

**Table S2:** microarray analysis of differentially regulated microRNAs at 5hpi.

| Column ID | p-value(Shigella-Infected-5h vs. NI) | Fold-Change(Shigella-Infected-5h vs. NI) |
| --- | --- | --- |
| MATURE_hsa-mir-6762-5p_at | 5,2266E-09 | 55,7357 |
| MATURE_hsa-mir-6797-5p_at | 2,2E-07 | 8,40346 |
| HAIRPIN_hsa-mir-6762_at | 1,2135E-06 | 2,13744 |
| MATURE_hsa-mir-4732-5p_at | 1,637E-05 | 5,28969 |
| MATURE_hsa-mir-6787-5p_at | 0,00004714 | 3,23986 |
| MATURE_hsa-mir-3687_at | 0,00028901 | 5,13259 |
| HAIRPIN_hsa-mir-3687-1_at | 0,00034538 | 5,17987 |
| HAIRPIN_hsa-mir-3687-2_at | 0,00034538 | 5,17987 |
| MATURE_hsa-mir-7150_at | 0,00036487 | 2,53181 |
| MATURE_hsa-mir-665_at | 0,00060187 | 2,77846 |
| MATURE_hsa-mir-371b-5p_at | 0,0007767 | 3,09814 |
| MATURE_hsa-mir-4505_at | 0,00106911 | 2,42365 |
| MATURE_hsa-mir-3937_at | 0,00154775 | 4,49102 |
| <u>MATURE_hsa-mir-4767_at</u> | 0,00170544 | -2,48664 |
| MATURE_hsa-mir-4726-5p_at | 0,00362434 | 2,02088 |
| MATURE_hsa-mir-4632-5p_at | 0,00631524 | 2,00295 |
| MATURE_hsa-mir-33b-3p_at | 0,00742774 | -2,39137 |
| MATURE_hsa-mir-4492_at | 0,00945298 | 2,1274 |
| <u>MATURE_hsa-mir-5100_at</u> | 0,0100979 | 3,08178 |
| MATURE_hsa-mir-663b_at | 0,0103003 | 2,2645 |
| MATURE_hsa-mir-3181_at | 0,0133326 | -2,34793 |
| MATURE_hsa-mir-1202_at | 0,0134169 | 2,40799 |
| <u>HAIRPIN_hsa-mir-5100_at</u> | 0,0136018 | 2,47264 |
| MATURE_hsa-mir-4649-5p_at | 0,01426 | 2,01001 |
| MATURE_hsa-mir-6795-5p_at | 0,0145134 | 2,22981 |
| MATURE_hsa-mir-1183_at | 0,0170046 | 2,22751 |
| MATURE_hsa-mir-370-3p_at | 0,0176671 | 2,377 |
| MATURE_hsa-mir-1234-3p_at | 0,0182928 | -2,60442 |
| MATURE_hsa-mir-6802-5p_at | 0,0232521 | 2,15321 |
| MATURE_hsa-mir-3197_at | 0,0237573 | 2,18409 |
| MATURE_hsa-mir-6075_at | 0,0302289 | 2,13437 |
| MATURE_hsa-mir-1247-5p_at | 0,0316243 | 2,12843 |
| MATURE_hsa-mir-4513_at | 0,0338213 | 2,00043 |
| MATURE_hsa-mir-6726-5p_at | 0,0364229 | 2,0802 |

**Table S3:**
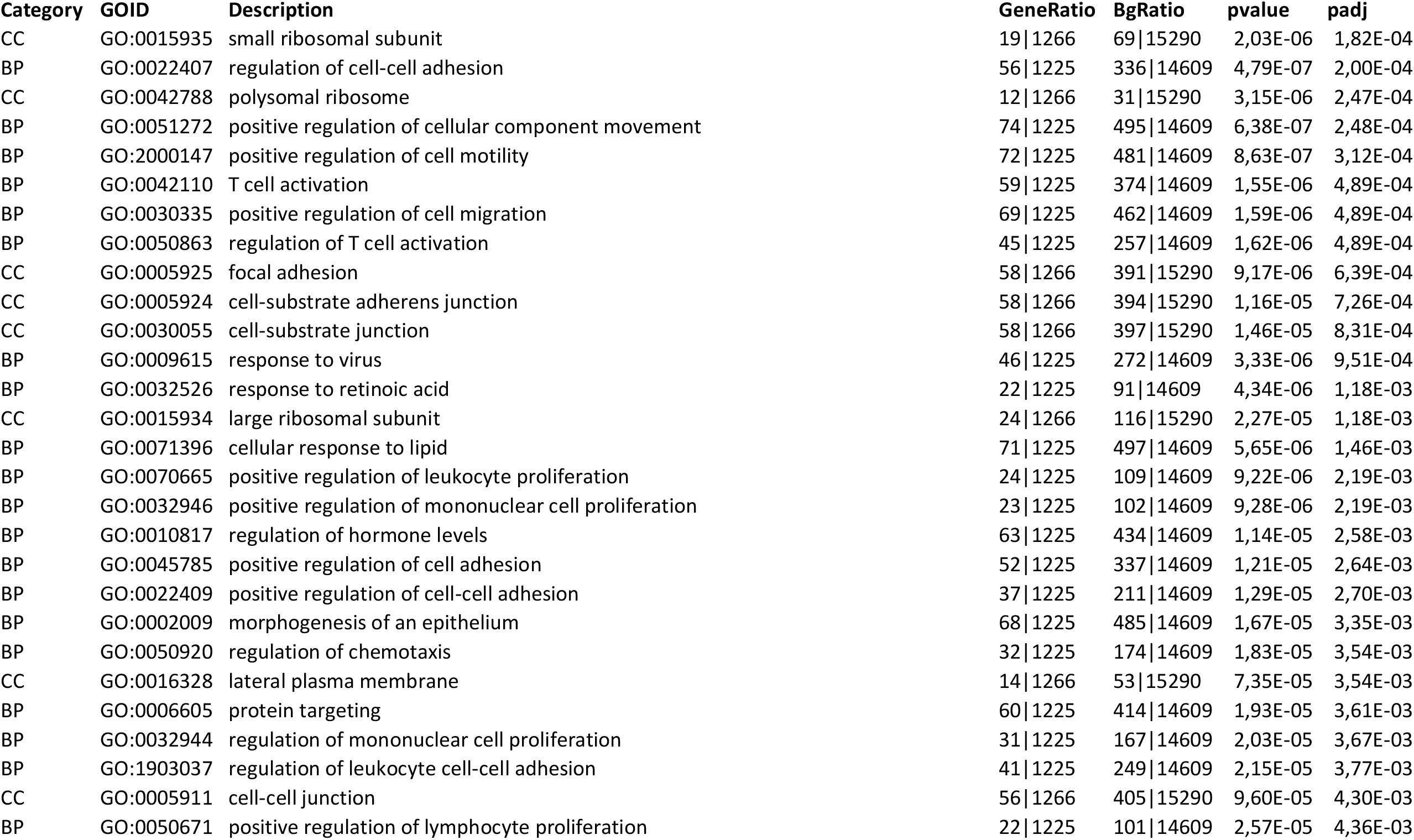
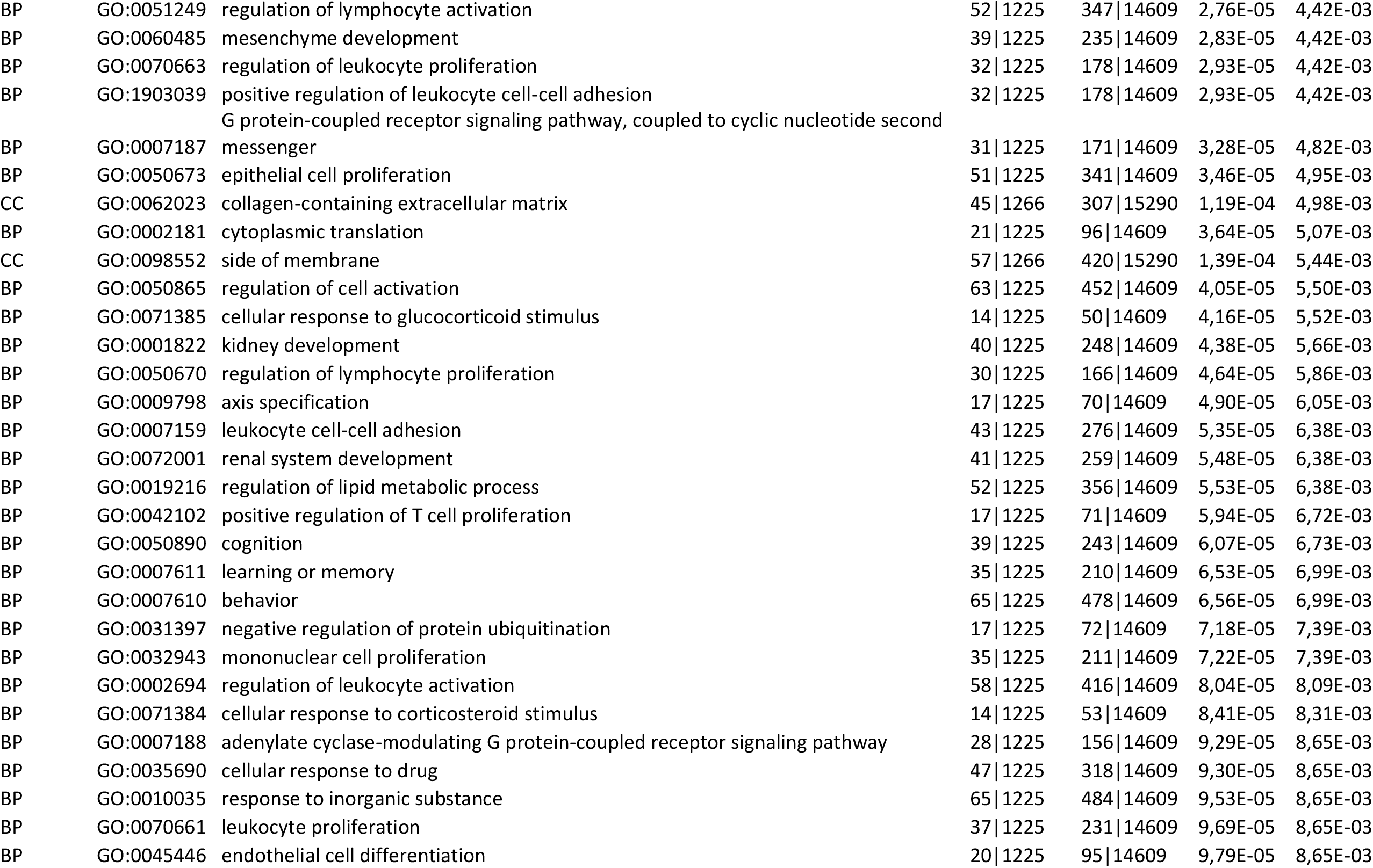
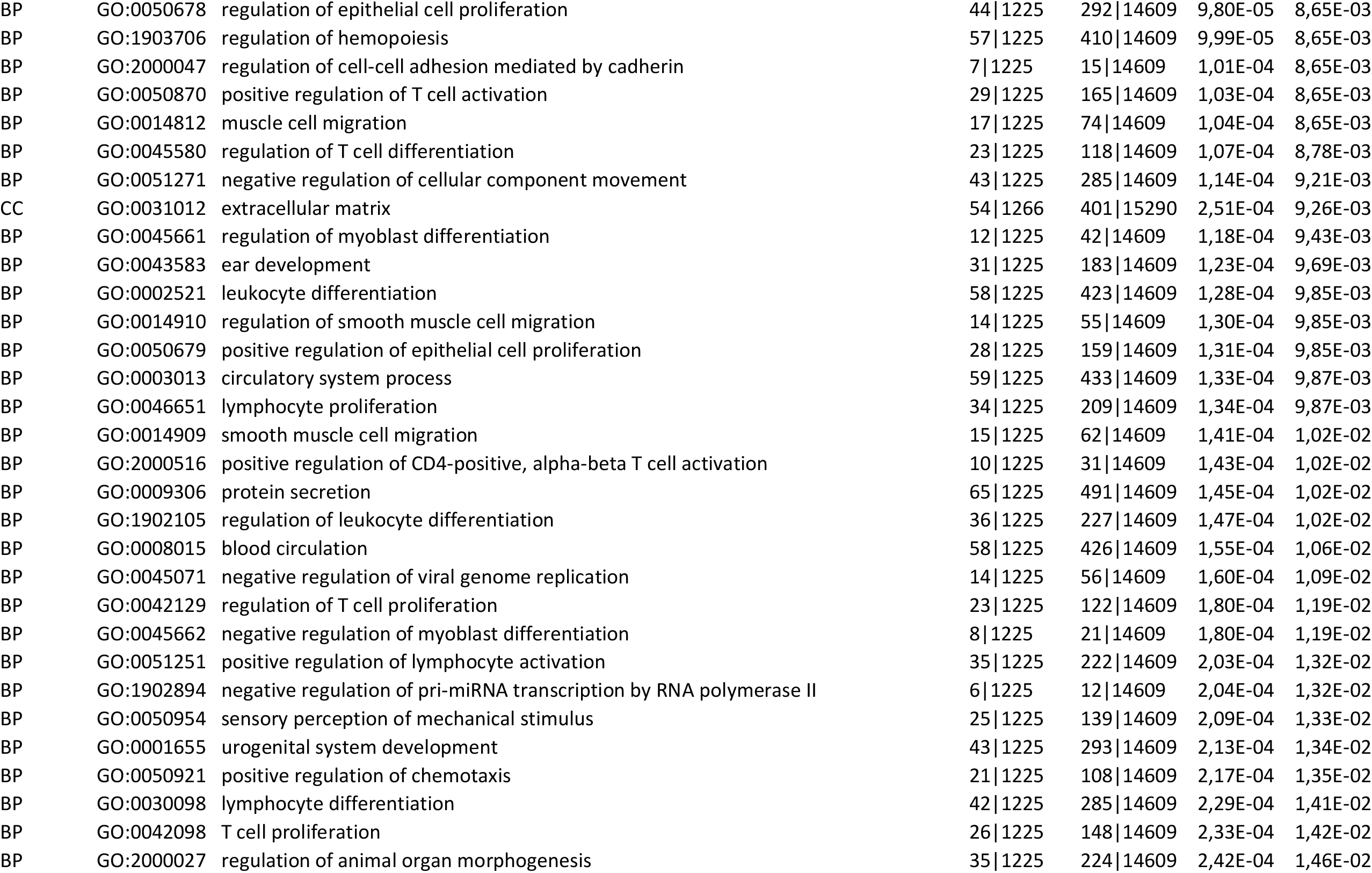
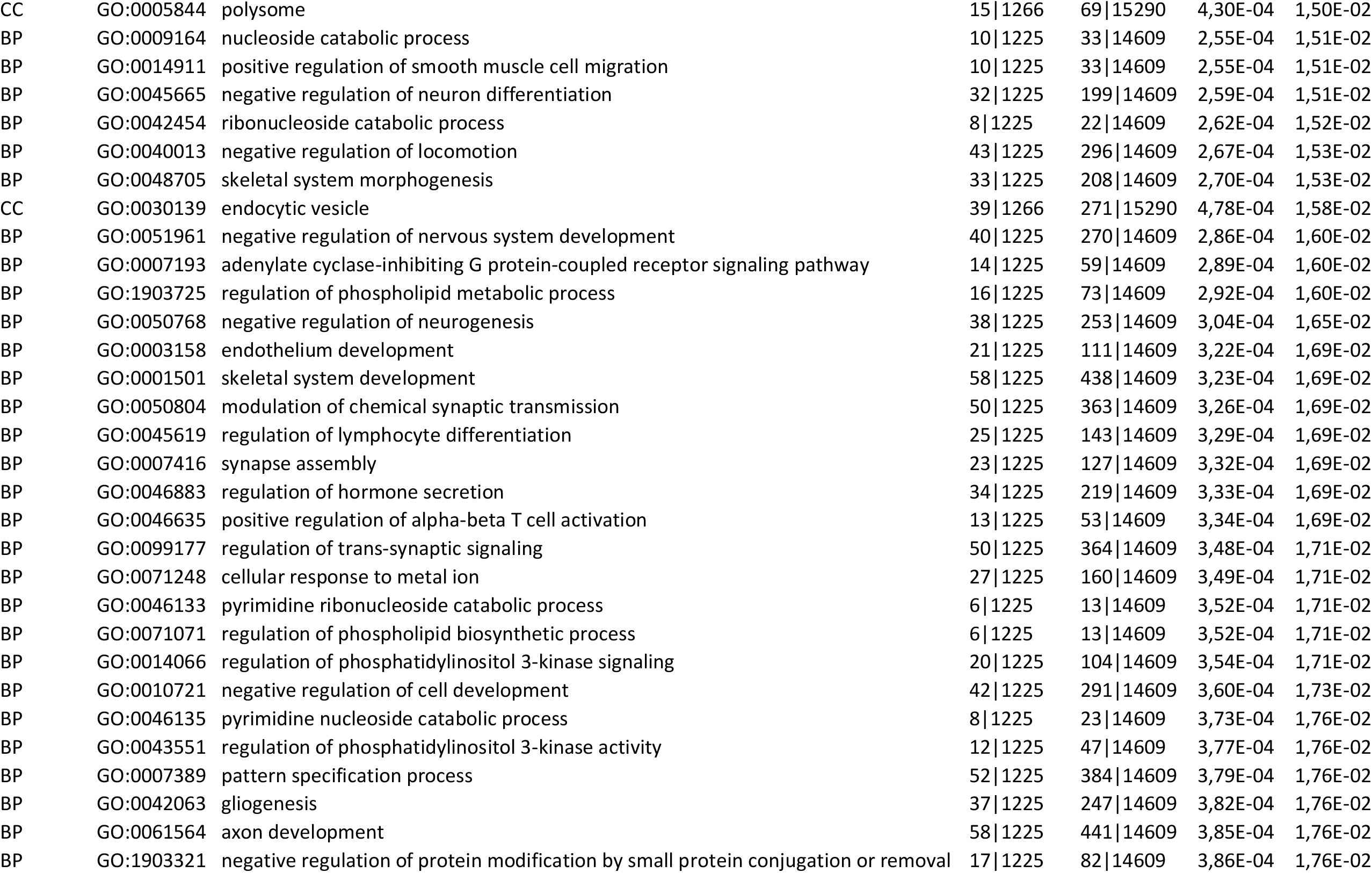
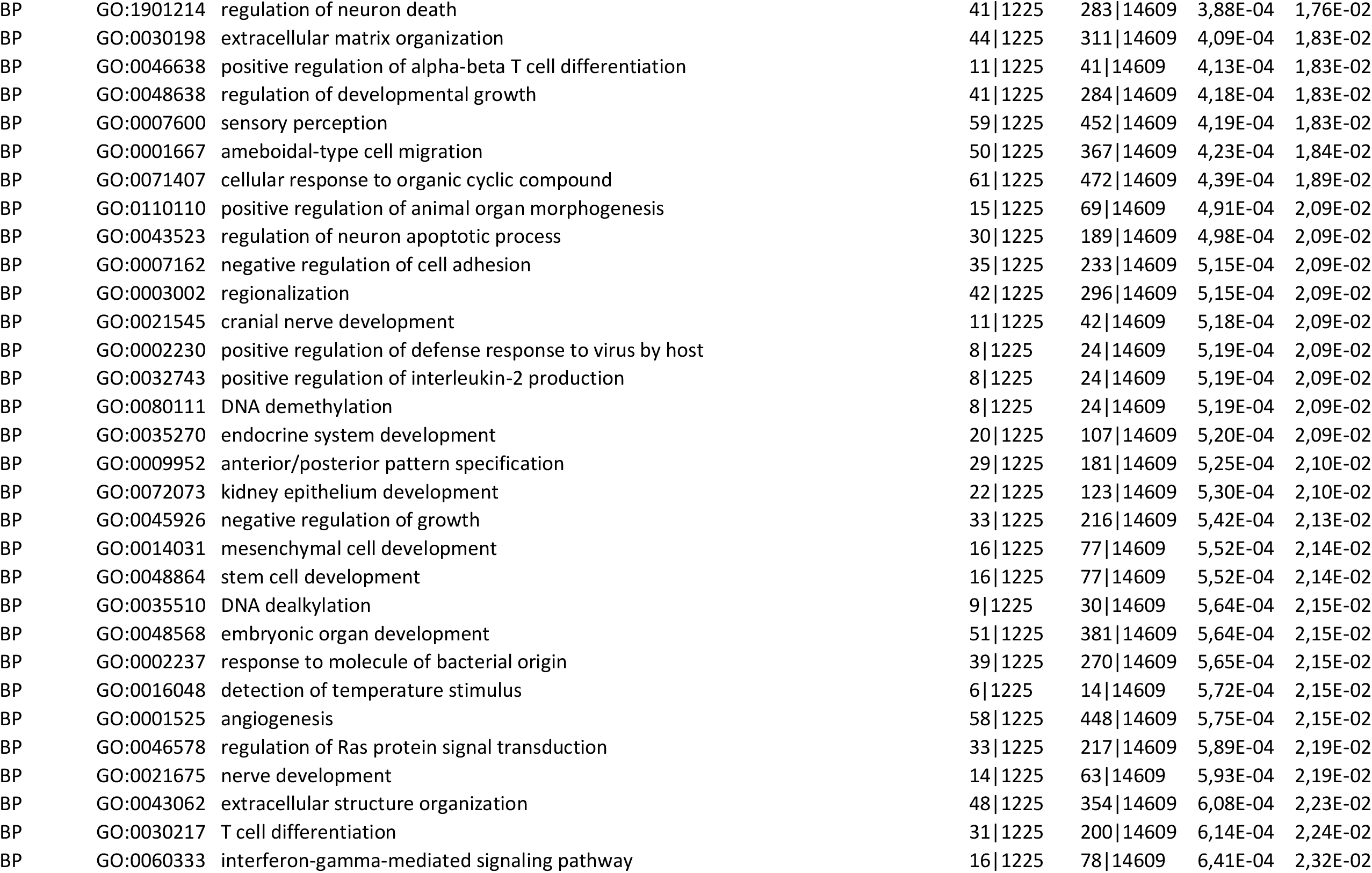
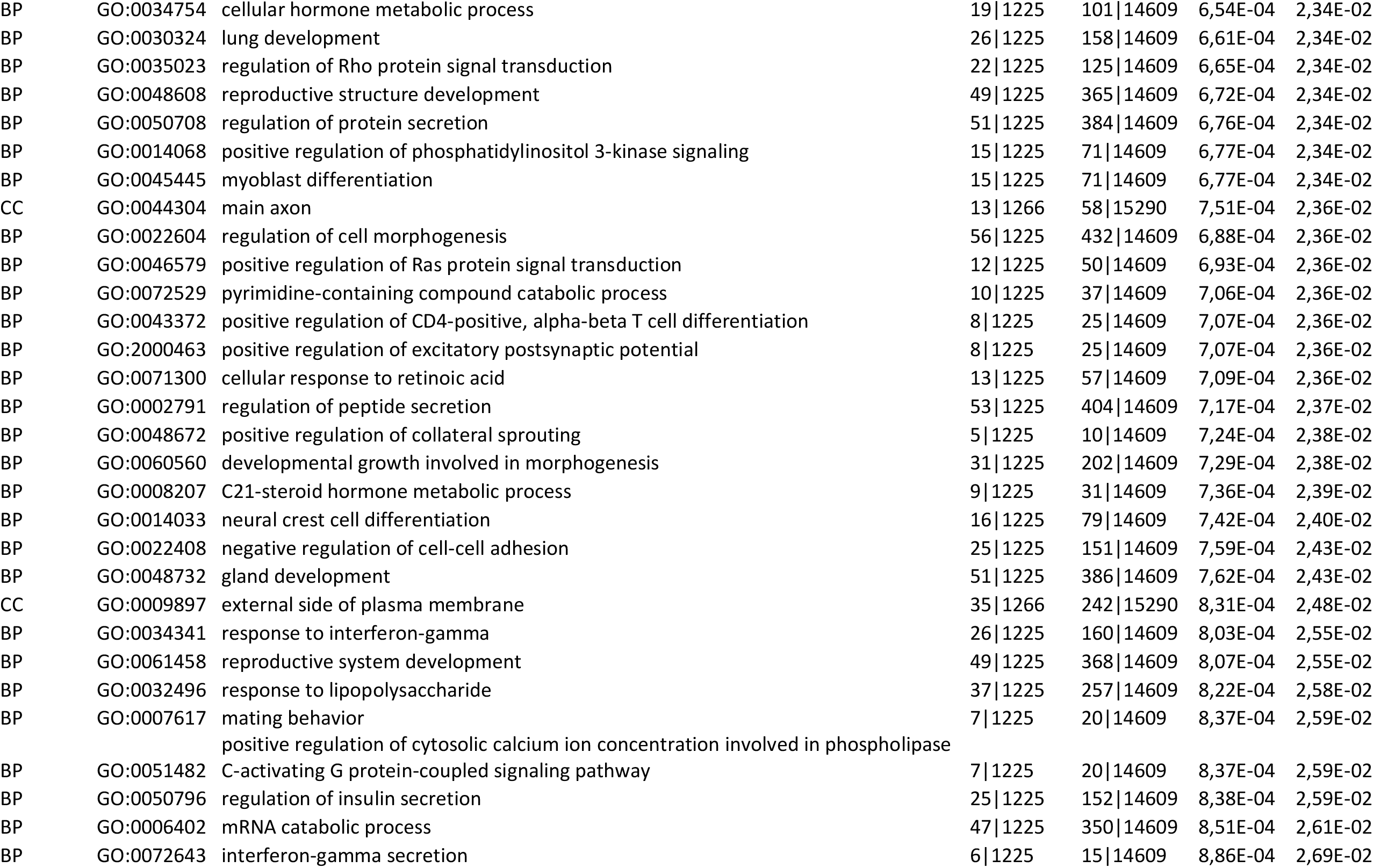
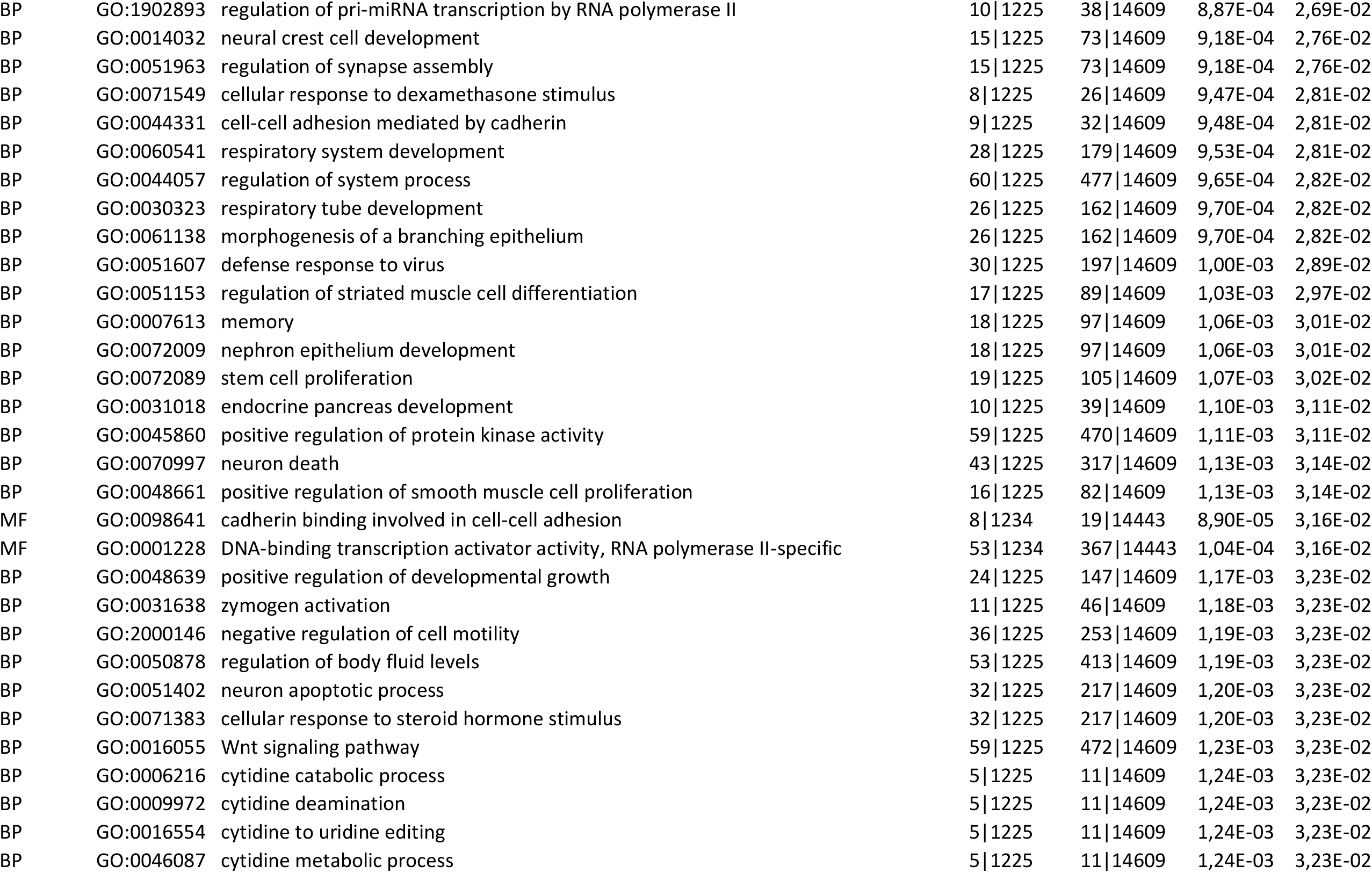
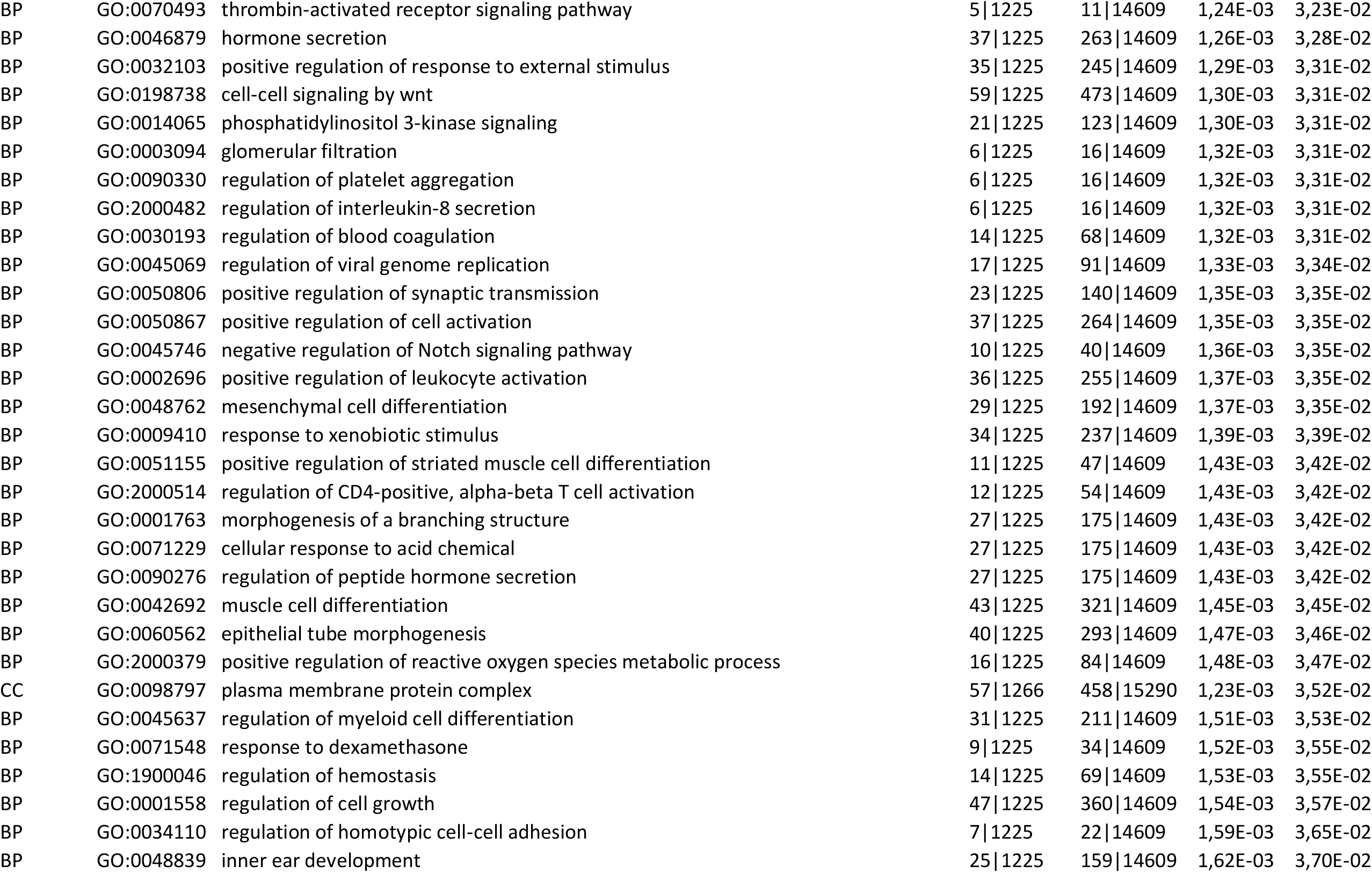
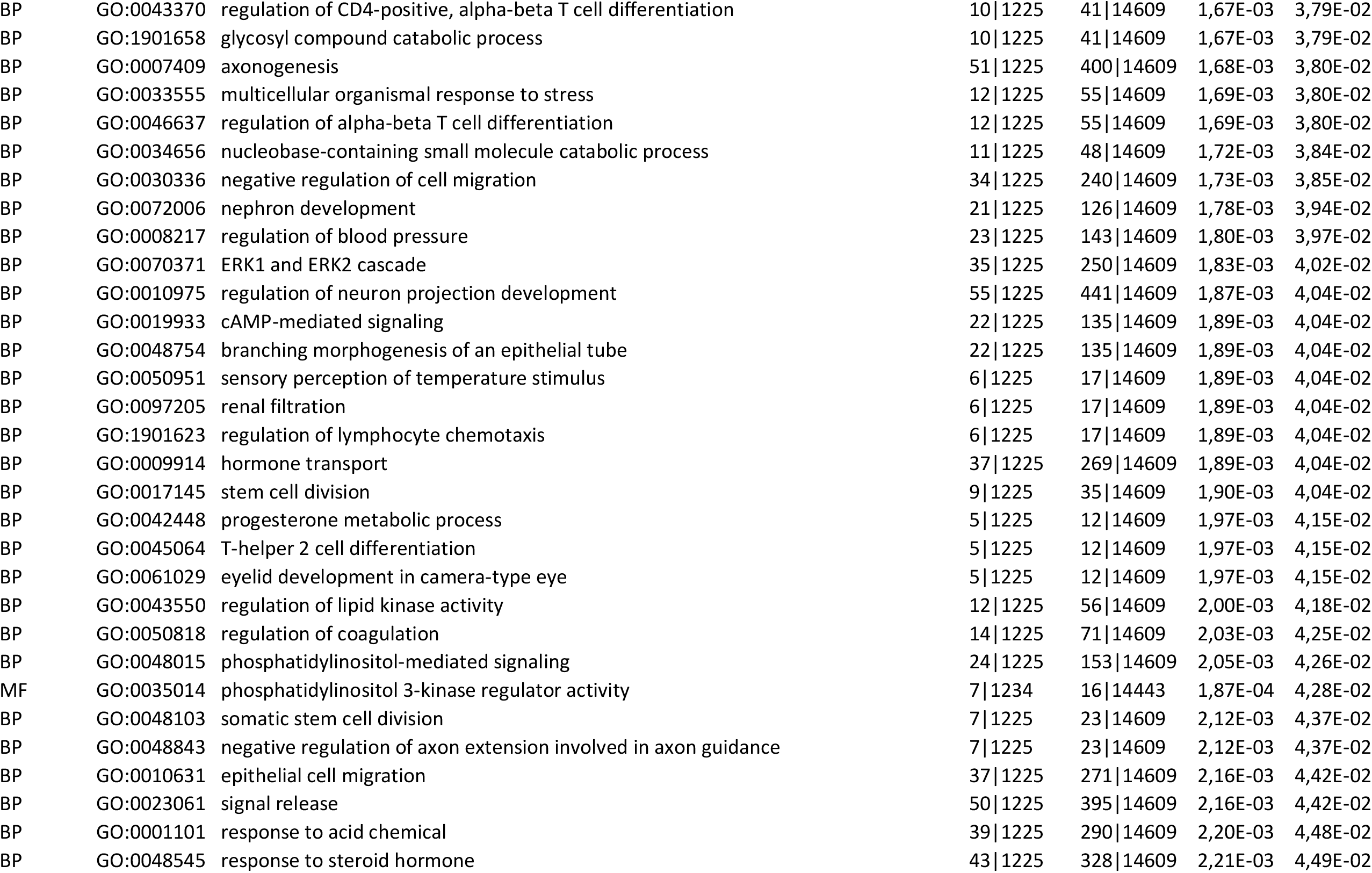
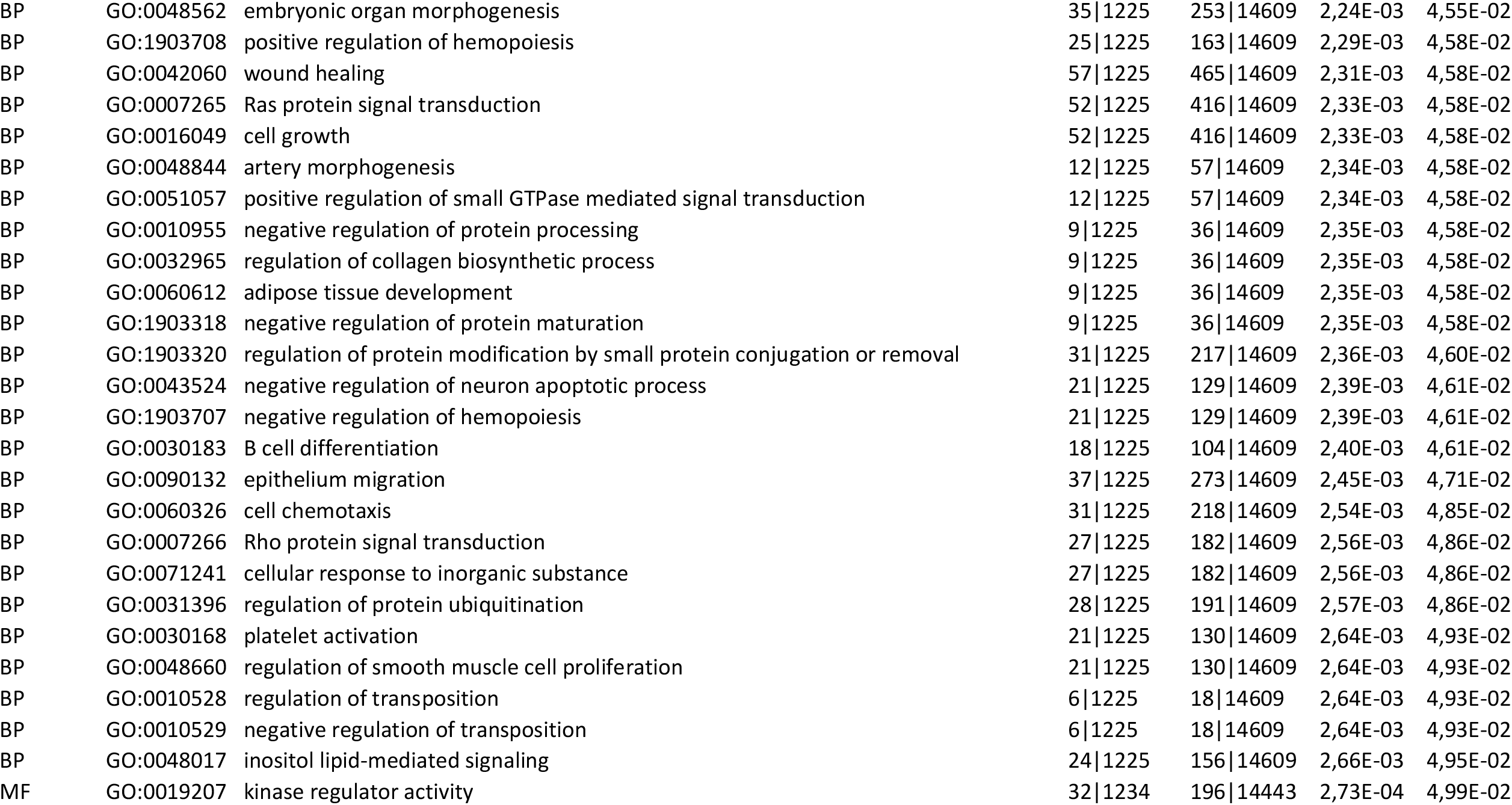
GO enrichement analysis - Upregulated pathways in HCT116 induced miR-6762-5p vs induced NT control (p<0.05)

**Table S4:** GO enrichement analysis - Downregulated pathways in HCT116 induced miR-6762-5p vs induced NT control (p<0.05)

| **Category** | **GOID** | **Description** | **GeneRatio** | **BgRatio** | **pvalue** | **padj** |
| --- | --- | --- | --- | --- | --- | --- |
| CC | GO:0031252 | cell leading edge | 52\|1132 | 375\|15290 | 8,13E-06 | 2,98E-03 |
| CC | GO:0030117 | membrane coat | 20\|1132 | 94\|15290 | 1,42E-05 | 2,98E-03 |
| CC | GO:0048475 | coated membrane | 20\|1132 | 94\|15290 | 1,42E-05 | 2,98E-03 |
| CC | GO:0030027 | lamellipodium | 31\|1132 | 187\|15290 | 1,81E-05 | 2,98E-03 |
| BP | GO:0030204 | chondroitin sulfate metabolic process | 11\|1103 | 35\|14609 | 3,32E-05 | 4,98E-02 |
| BP | GO:0006914 | autophagy | 60\|1103 | 472\|14609 | 4,47E-05 | 4,98E-02 |
| BP | GO:0061919 | process utilizing autophagic mechanism | 60\|1103 | 472\|14609 | 4,47E-05 | 4,98E-02 |
| BP | GO:0061572 | actin filament bundle organization | 25\|1103 | 142\|14609 | 5,64E-05 | 4,98E-02 |
| BP | GO:0030641 | regulation of cellular pH | 18\|1103 | 86\|14609 | 6,13E-05 | 4,98E-02 |
| BP | GO:0051452 | intracellular pH reduction | 13\|1103 | 50\|14609 | 6,14E-05 | 4,98E-02 |
| BP | GO:0043087 | regulation of GTPase activity | 55\|1103 | 429\|14609 | 7,32E-05 | 4,98E-02 |
| BP | GO:0045851 | pH reduction | 13\|1103 | 51\|14609 | 7,67E-05 | 4,98E-02 |

**Table S5:**
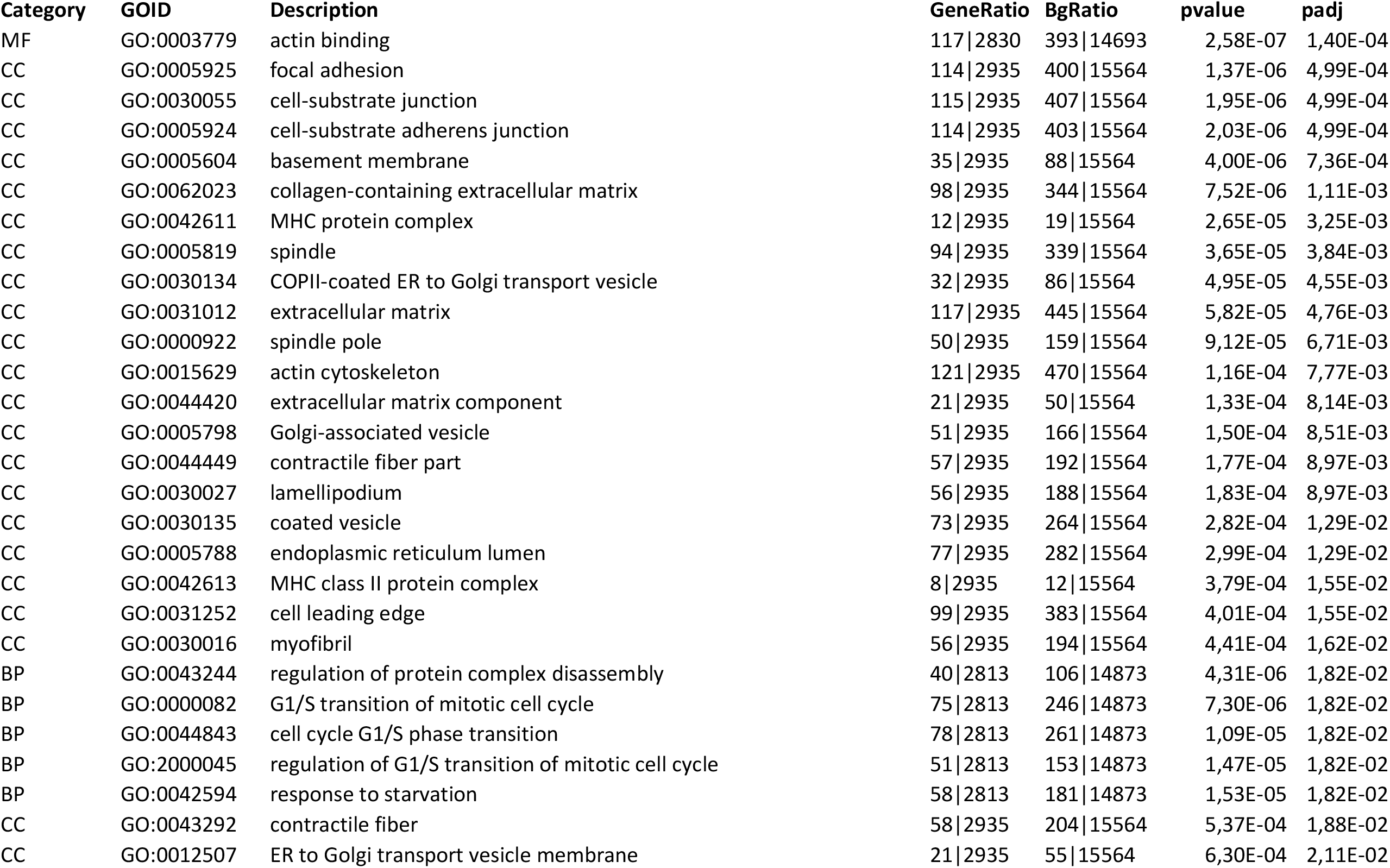
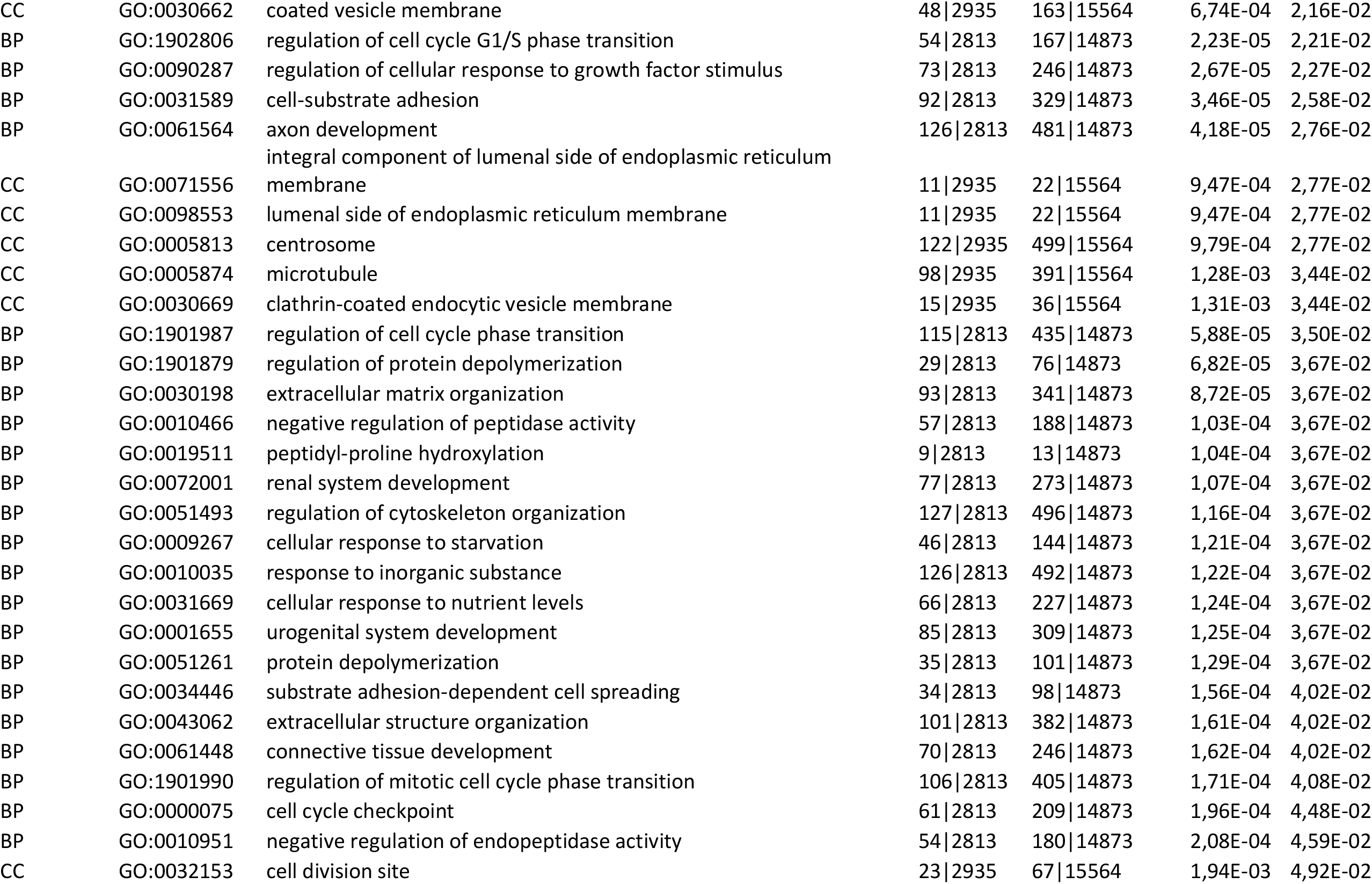
GO enrichement analysis - Upregulated pathways in HeLa induced miR-6762-5p vs induced NT control (p<0.05)

**Table S6:** GO enrichement analysis - Downregulated pathways in HeLa induced miR-6762-5p vs induced NT control (p<0.05)

| Category | GOID | Description | GeneRatio | BgRatio | pvalue | padj |
| --- | --- | --- | --- | --- | --- | --- |
| BP | GO:0050804 | modulation of chemical synaptic transmission | 127\|3299 | 382\|14873 | 3,16E-07 | 1,00E-04 |
| BP | GO:0051962 | positive regulation of nervous system development | 152\|3299 | 479\|14873 | 5,80E-07 | 1,75E-04 |
| CC | GO:0097060 | synaptic membrane | 118\|3428 | 365\|15564 | 2,75E-06 | 2,07E-04 |
| CC | GO:0005840 | ribosome | 79\|3428 | 224\|15564 | 3,51E-06 | 2,40E-04 |
| CC | GO:0043025 | neuronal cell body | 134\|3428 | 434\|15564 | 8,88E-06 | 5,37E-04 |
| CC | GO:0015934 | large ribosomal subunit | 47\|3428 | 118\|15564 | 9,26E-06 | 5,37E-04 |
| CC | GO:0098978 | glutamatergic synapse | 102\|3428 | 315\|15564 | 1,18E-05 | 6,35E-04 |
| CC | GO:0044297 | cell body | 149\|3428 | 495\|15564 | 1,30E-05 | 6,55E-04 |
| CC | GO:0043197 | dendritic spine | 57\|3428 | 156\|15564 | 2,40E-05 | 1,13E-03 |
| CC | GO:0043235 | receptor complex | 109\|3428 | 347\|15564 | 2,66E-05 | 1,18E-03 |
| CC | GO:0044309 | neuron spine | 57\|3428 | 158\|15564 | 3,66E-05 | 1,53E-03 |
| BP | GO:0031345 | negative regulation of cell projection organization | 65\|3299 | 176\|14873 | 5,64E-06 | 1,62E-03 |
| CC | GO:0033267 | axon part | 106\|3428 | 340\|15564 | 4,81E-05 | 1,91E-03 |
| CC | GO:0005911 | cell-cell junction | 126\|3428 | 420\|15564 | 6,98E-05 | 2,63E-03 |
| CC | GO:0045177 | apical part of cell | 101\|3428 | 327\|15564 | 1,07E-04 | 3,83E-03 |
| CC | GO:0150034 | distal axon | 80\|3428 | 249\|15564 | 1,32E-04 | 4,46E-03 |
| CC | GO:0030133 | transport vesicle | 107\|3428 | 352\|15564 | 1,36E-04 | 4,46E-03 |
| CC | GO:0070382 | exocytic vesicle | 64\|3428 | 191\|15564 | 1,60E-04 | 4,87E-03 |
| CC | GO:0048786 | presynaptic active zone | 28\|3428 | 66\|15564 | 1,61E-04 | 4,87E-03 |

**Table S7:**
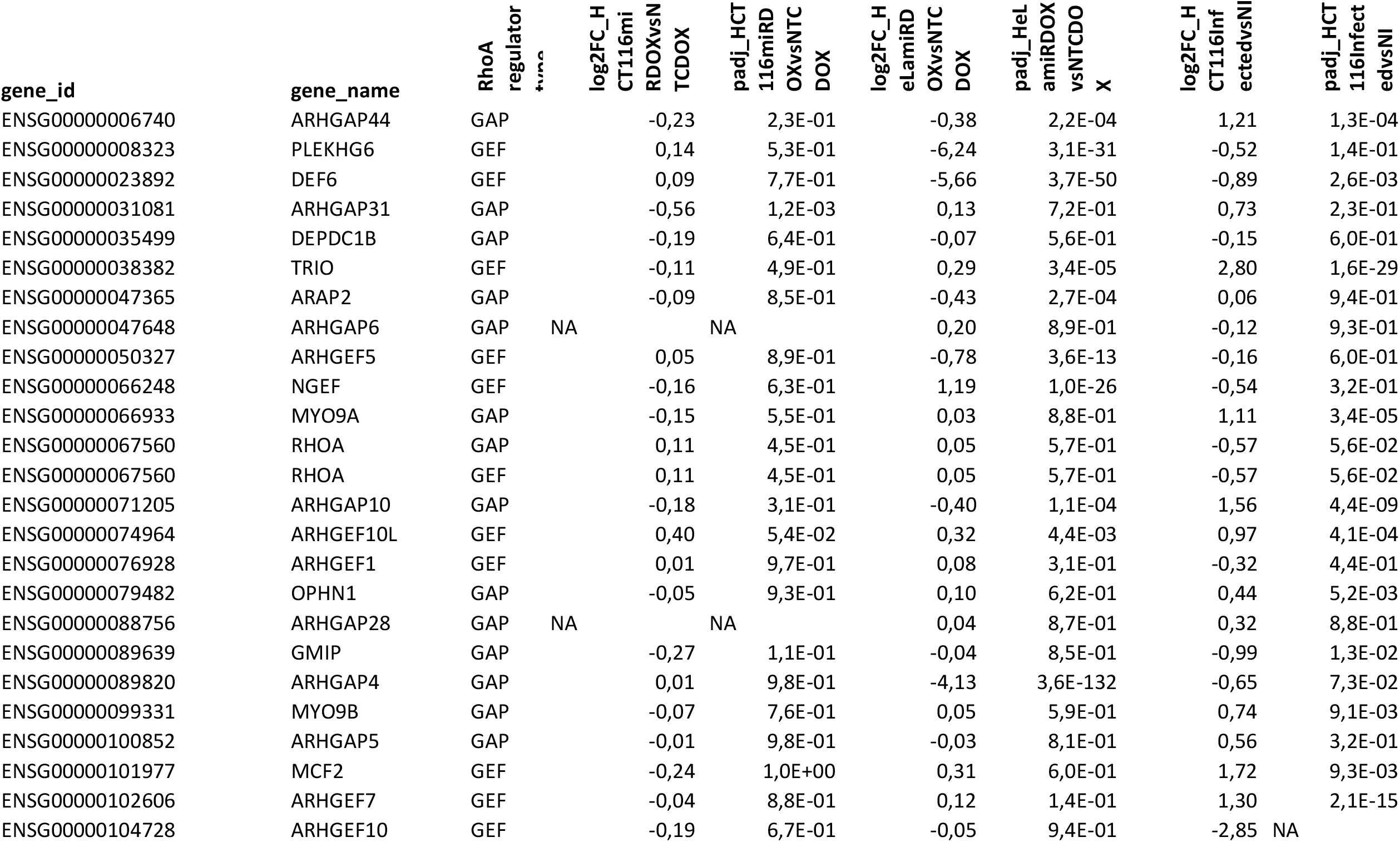
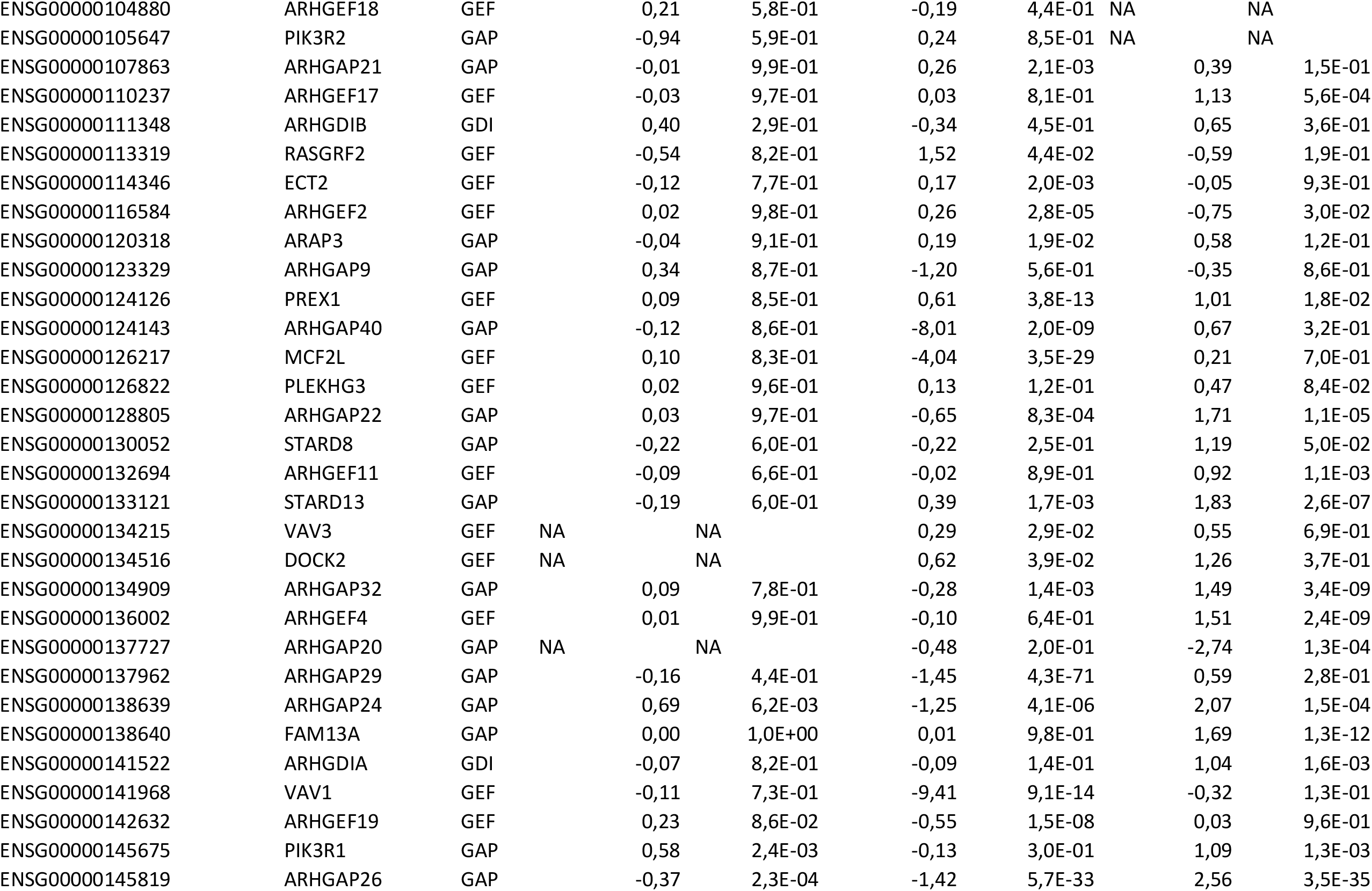
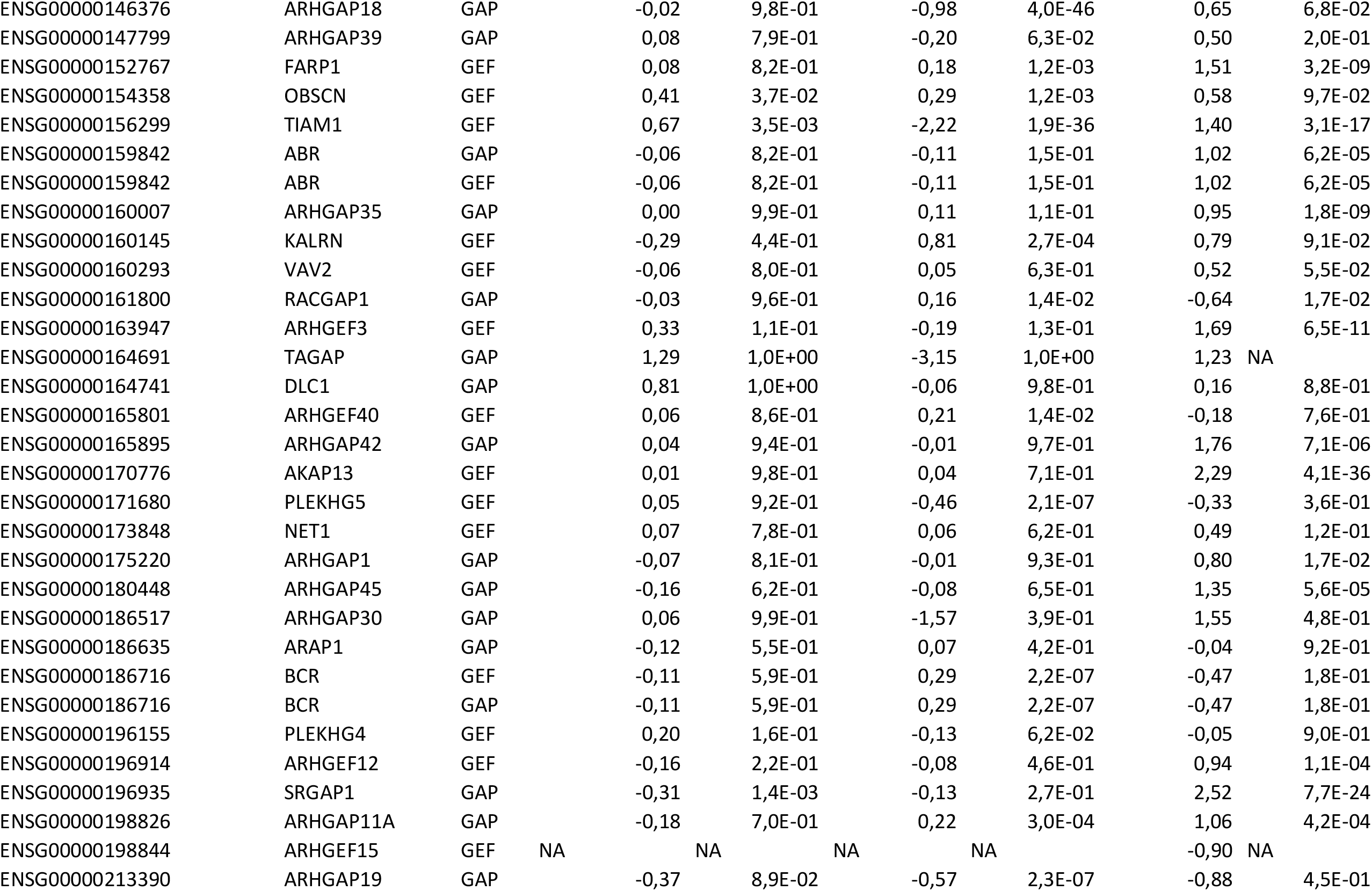
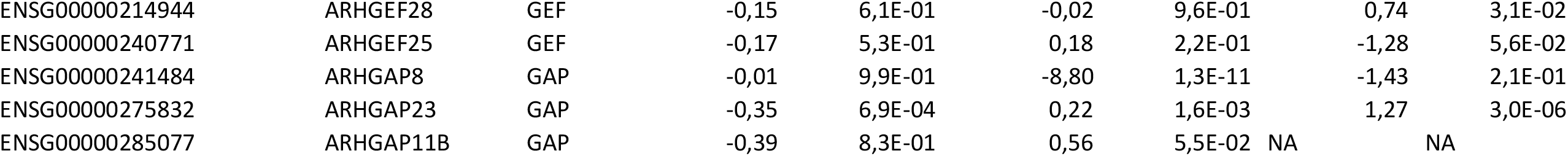
Expression of RhoA GEF, GAP and GDI in HCT116^miR^ vs. HCT116^NTC^, HeLa^miR^ vs. HeLa^NTC^, and *Shigella*-infected HCT116 vs. non-infected.

## Material and Methods

### Cell lines, bacterial strain, and culture reagents

*Shigella flexneri* serotype 5 strain M90T was grown in trypticase soy broth. The same strain was used carrying a pSU18-mCherry plasmid for plaque assays. HeLa cells were from ATCC (CCL-2). HCT116 cells were a gift from B. Vogelstein (Johns Hopkins University, Baltimore, MD, USA). HeLa and HCT116 cells were cultivated in DMEM GlutaMAX (Life Technologies, 10566016) supplemented with MEM non-essential amino acids (Life Technologies, 11140050), 1mM sodium pyruvate (Life Technologies, 11360070), 100U/ml Penicillin-Streptomycin (Life Technologies, 15140122), and 10% fetal calf serum (Eurobio, CVFSVF00). When indicated, cells were treated for an hour with the ROCK inhibitor Y27632 at a concentration of 15µM in DMSO. The same volume of DMSO was added to negative control samples.

### Inducible microRNA stable cell line generation

HeLa and HCT116 cell lines were transduced with shMIMIC Inducible Lentiviral microRNAs - vector piSMART mCMV/TurboGFP with mimic sequence 5’-CGGGGCCAUGGAGCAGCCUGUGU-3’ for miR-6762-5p (Horizon Discovery, VSH6904-224674299) and sequence 5’-CACACAACAUCUAAACCAGGGA-3’ for the non-targeting control (Horizon Discovery, VSC6570). Transduction was performed 24h after cell plating on 40% cell confluence with 8µg/ml polybrene in previously described cell medium without antibiotics at MOI 0.1 of lentiviral particles. To minimize expression leakage of transgene we cultured cells with fetal calf serum without tetracycline (Eurobio, CVFSVF1101). Complete medium was added 20h after transduction. Medium was changed 48h after transduction. 72h after transduction cells were expanded and selected in 0,5µg/ml puromycine for four days. Doxycyline induction of microRNA expression was performed by adding 250ng of doxycycline to the cell medium.

### MicroRNA inhibitor transfection

miRIDIAN microRNA hairpin inhibitor for human miR-6762-5p and negative control #1 were purchased from Horizon Discovery (IH-302910-01 and IN-001005-01). microRNA inhibitors were transfected by inverse transfection at a 50nM concentration with DharmaFECT 1 transfection reagent (Horizon Discovery, T-2001) in Opti-MEM (Life Technologies, 11058021) in the absence of antibiotics.

### Bacterial infection

Overnight cultures grown at 30°C with shaking 160rpm were diluted 1:35 and grown at 37°C with shaking at 160rpm until OD = 0.7 was reached. Bacterial pellet was resuspended in DMEM without serum and bacteria suspension was added to 30 min serum - starved and 70% confluent cells. A multiplicity of infection of 100 was used, except when otherwise mentioned. Plates were centrifuged for 10min at 2000rpm at room temperature. Plates were then incubated for 30 min in humid atmosphere, 37°C temperature and 5% CO2. Cells were then washed three times with PBS. Then, DMEM without serum, and supplemented with 50 µg/ml Gentamicin, was added to the cells. Time 0 of infection was counted from here.

### Plaque assay

MicroRNA inducible cell lines were induced for 3 days prior to infection. Bacterial suspension in DMEM without serum was added to 30 min serum-starved cells. Bacterial infection was caried out on 100% confluent cells at a multiplicity of infection of 0.01. Cells with bacterial suspension were incubated for 2h in a humid atmosphere, 37°C temperature and 5% CO2. Cells were then washed three times with PBS. Then, DMEM with 10% fetal calf serum, and supplemented with 50 µg/ml Gentamicin, was added to the cells. Plates were incubated for 24h in a humid atmosphere, 37°C temperature and 5% CO2. Cells were then fixed and stained with DAPI for imaging.

### RNA extraction

Total RNA was purified by TRIzol (Life Technologies, 15596026) using 1ml of reagent for each 6 million of cells or less, according to the manufacturer’s instructions. Briefly, cells were lysed for 5 min by addition of the TRIzol into the cell culture dish. 200µl of chloroform (Sigma-Aldrich, 372978) was added for phase separation. After 15min centrifugation at 12,000g and 4°C, the upper phase was collected, and an equal volume of isopropanol (Sigma-Aldrich, I9516) was added. After homogenization, the samples were left at −20°C overnight for precipitation. Total RNA was precipitated by centrifugation at 20,000g for >30min at 4°C. The pellet was then washed twice with ice-cold 70% nuclease-free ethanol (Sigma-Aldrich, 51976). Finally, the pellet was shortly left to dry at room temperature and then resuspended in nuclease-free H_2_O (Life Technologies, 10977035) and kept at −80°C between uses.

### Microarray analysis of microRNA expressions

The total RNAs from 5h *Shigella*-infected and non-infected control HCT116 cells were collected. Three independent experiments were performed and analyzed. Microarray microRNA expression quantification was carried out at the Genom’ic Core Facility at the Institut Cochin, University of Paris, using the GeneChip™ miRNA 4.0 Array (ThermoFisher Scientific, 902411) according to the manufacturer’s instruction. Relative expression and statistical analysis were performed using the Transcriptome Analysis Console (TAC) Software (ThermoFisher Scientific). Differentially expressed microRNAs were defined as microRNAs with a higher than a 2-fold-change expression difference and an associated adjusted p-value inferior to 0.05.

### RNA sequencing

HCT116^NTC^, HCT116^miR^, HeLa^NTC^, and HeLa^miR^ cells were stimulated with 100ng/ml doxycycline for 48h to induce microRNA expression. Three independent experiments were performed. Library, sequencing and bioinformatic analysis was performed by Novogene Co., Ltd as follows. For library preparation, a total amount of 1 μg RNA per sample was used as input material. Sequencing libraries were generated using NEBNext® UltraTM RNA Library Prep Kit for Illumina® (NEB, USA) following manufacturer’s recommendations and index codes were added to attribute sequences to each sample. To select cDNA fragments of preferentially 250∼300 bp in length, the library fragments were purified with AMPure XP system (Beckman Coulter, Beverly, USA) and library quality was assessed on the Agilent Bioanalyzer 2100 system. The library preparations were sequenced on an Illumina platform and paired-end reads were generated. Raw reads were first processed through fastp. In this step, clean reads were obtained by removing reads containing adapter and poly-N sequences and reads with low quality from raw data. At the same time, Q20, Q30 and GC content of the clean data were calculated. All the downstream analyses were based on the clean data with high quality. Human reference genome Hg38 and gene model annotation files were downloaded from genome website browser (NCBI/UCSC/Ensembl) directly. Indexes of the reference genome was built using STAR and paired-end clean reads were aligned to the reference genome using STAR (v2.5). HTSeq v0.6.1 was used to count the read numbers mapped of each gene. And then FPKM of each gene was calculated. Differential expression analysis between two groups was performed using the DESeq2 R package (2_1.6.3). The resulting P-values were adjusted using the Benjamini and Hochberg’s approach for controlling the False Discovery Rate (FDR). Genes with an adjusted P-value <0.05 found by DESeq2 were considered differentially expressed. Finally, Gene Ontology (GO) enrichment analysis of differentially expressed genes was implemented by the clusterProfiler R package, in which gene length bias was corrected. GO terms with corrected P-value less than 0.05 were considered significantly enriched by differential expressed genes.

### Reverse transcription and quantitative PCR

Messenger RNAs were reverse transcribed using the RevertAid H Minus First Strand cDNA Synthesis Kit (Life Technologies, K1632) using oligo-dT primers. Quantitative PCR was performed using the Brilliant III Ultra-Fast SYBR Green qRT-PCR Master Mix (Agilent, 600886) on a Mx3005P Real-Time PCR System (Agilent). The primers used for cDNA amplification were the following: RLP0 forward 5’-AGGTGTTCGACAATGGCAGCAT-3’ and reverse 5’-TGCAGACAGACACTGGCAACAT-3’; MYO1E forward 5’-CCAGAACTCCAGCAGTTCGT-3’ and reverse 5’-AGGTCTCGCTTTACACCCTTG-3’; MYH15 forward 5’-GGAGTGAGAGTGGCGAGTTC-3’ and reverse 5’-AGAAGGTCACAGTCACGCTG-3’; FBLIM1 forward 5’-GAACCCTGCTACCAGGACAC-3’ and reverse 5’-TCTTTCCCATCCCGAGGGAT-3’; TMSB4X forward 5’-TGCCTTCCAAAGAAACGATTGA-3’ and reverse 5’-CCTGCCAGCCAGATAGATAGAC-3’.

For microRNAs, the stem-loop RT-qPCR technique was used. The protocol from Varkonyi-Gasic and Hellens (2011) was used for reverse transcription. Briefly, 10ng total RNA was reverse-transcribed for each microRNA in a 20µl mix containing 250µM dNTP mix, 10mM DTT, 50U SuperScript III reverse transcriptase, 1x First Strand buffer (Life Technologies, 18080044), 4U RNases inhibitor (Life Technologies, 10777019), 50µM denatured stem-loop oligonucleotide. The reverse transcription was performed using pulsed RT protocol as follows: 30min incubation at 16°C, followed by 60 cycles of – 30 seconds 30°C, 30 seconds 42°C and 1 second 50°C – finished by an inactivation step of 5 minutes at 85°C. The following stem-loop oligonucleotides were used for: miR-103a-3p 5’-GTTGGCTCTGGTGCAGGGTCCGAGGTATTCGCACCAGAGCCAACTCATAG-3’; miR-6762-5p 5’-GTTGGCTCTGGTGCAGGGTCCGAGGTATTCGCACCAGAGCCAACACACAG-3’; for the non-targeting control microRNA 5’-GTTGGCTCTGGTGCAGGGTCCGAGGTATTCGCACCAGAGCCAACTCCCTG-3’. The quantitative PCR was performed using the Brilliant III Ultra-Fast SYBR Green qRT-PCR Master Mix (Agilent, 600886) on a Mx3005P Real-Time PCR System (Agilent). The 20µl reaction was as follows: 1X SYBR green master mix, 500µM of forward and reverse primers, and 2µl of the reverse transcription. Cycling conditions were as follows: 3min et 95°C, followed by 40 cycles of 20 seconds at 95°C, 20 seconds at 60°C, and fluorescence acquisition. PCR was followed by melting curve analysis to ensure amplification of a single fragment of the expected melting temperature. PCR primers were the following: miR-103a-3p forward 5’-TGTTTTTTTGAGCAGCATTGTACAG-3’, miR-6762-5p forward 5’-TTGTTCGGGGCCATGGAG-3’, non-targeting control microRNA forward 5’-GTTTGGCACACAACATGTAAAC-3’, and universal reverse 5’-GTGCAGGGTCCGAGGT-3’.

### GTPases activation assay

For the analysis of small GTPases activation, we used the RhoA/Rac1/Cdc42 Activation Assay Combo Biochem Kit (Cytoskeleton, Inc., BK030). The pull-down was performed following manufacturer’s instructions.

### Protein relative quantification by Western blot

Proteins were collected in cell lysis buffer (50mM Tris pH 7.5, 10mM MgCl_2_, 0.5M NaCl, 2% Igepal) supplemented with 1X Protease inhibitor cocktail (Cytoskeleton, PIC02). Lysates were cleared by 1min centrifugation at 10,000g at 4°C and the collected supernatants were snap-frozen in liquid nitrogen. Samples were stored at −80°C until analyzed. 50µg of lysate were denatured by addition of Leammli sample buffer supplemented with β-mercaptoethanol and by incubating for 5 min at 95°C with shaking. Proteins were separated on a 1.5mm thick, 12% Tris/Glycine SDS-PAGE. Liquid transfer was performed on Immobilon-P Membrane, PVDF, 0.45 µm membranes (Millipore, IPVH85R) in a 15% Methanol/Tris/Glycine buffer for 1h15min, 90V at 4°C. After transfer, membrane was rinsed in TBS buffer and let to dry for 20min. The membrane was then rehydrated with TBS-Tween 20 (0.05%) for 30min. Blocking was performed for 30min in 5% powdered skimmed milk suspended in TBST. Following a quick rinse in TBST, the primary antibody was incubated overnight at 4°C with shaking, resuspended in TBST. Anti-RhoA, anti-Rac1 and anti-CDC42 antibodies were used from the pull-down kit mentioned above at the recommended dilution. The anti-GAPDH antibody (Cell Signaling, #2118) was diluted 1:1000 in 5% bovine serum albumin in TBST. The anti-γTubulin antibody (Life Technologies, #MA1-850) was diluted 1:1000 in 5% bovine serum albumin (BSA) in TBST. Finally, the anti-TPCN1 antibody was generated against the C-terminal amino acid sequence of TPCN1 as previously described (Castonguay at al., 2017). Batch #3526 was used at a concentration of 1:1000.

### Phalloidin cell staining

Cells were cultivated on glass coverslips. Cell fixation was performed by incubating the coverslips in 3.2% paraformaldehyde in PBS for 20min at room temperature. Permeabilization was performed for 10min with 0.1% Triton-X 100 in PBS. Phalloidin-iFluor™ 647 Conjugate (AAT Bioquest, # 23127) and DAPI 1mg/ml stock (Roche, 10236276001) staining were performed simultaneously diluted 1:1000 in 2% BSA/PBS and incubated for 1h at room temperature. Coverslips were then mounted onto glass slides using ProLong™ Gold Antifade Mountant (Life Technologies, P36930).

### Microscopy image acquisition and quantification

Microscopy Images were acquired on a structured illumination fluorescent microscope Apotome 2 (Zeiss). Fluorescence quantification was performed using ImageJ software (Rasband, 1997-2018). In particular, stress fiber counting was performed as previously described (Wei et al., 2011) by counting the phalloidin fluorescence peaks over 1000 in intensity across a line spanning the width of the cell.

### miR-6762-5p sequence alignment

The comparative alignment of *MIR6762* human gene from the GRCh38 assembly with representative genomes of apes was done using the genomic alignment feature of the Ensembl database (Howe et al., 2021).

### *In sillico* analysis of RhoA activation modulators

The miRWalk2.0 database (Dweep and Gretz, 2015) was used to identify transcripts presenting putative 3’UTR target sites for miR-6762-5p. The miRWalk2.0 database collects the data from 4 different algorithms computing targeting potential of a miRNA on a transcript. We selected genes for which at least 2 algorithms detected a target site. RhoGAPs, RhoGEFs and RhoGDIs gene names were collected from the Reactome database (Jassal et al., 2020) categories: R-HSA-8980691; R-HSA-8981637; R-HSA-9012835. The list of miR-6762-5p putative targets was then crossed with the list of RhoA activation modulators.

### Statistical analysis

Statistical analysis were performed using GraphPad Prism 8.0.1.

